# Proteomic Assessment of SKBR3/HER2+ Breast Cancer Cellular Response to Lapatinib and Investigational Ipatasertib Kinase Inhibitors

**DOI:** 10.1101/2024.04.02.587656

**Authors:** Arba Karcini, Nicole R. Mercier, Iulia M. Lazar

## Abstract

Modern cancer treatment approaches aim at achieving cancer remission by using targeted and personalized therapies, as well as harnessing the power of the immune system to recognize and eliminate the cancer cells. To overcome a relatively short-lived response due to the development of resistance to the administered drugs, combination therapies have been pursued, as well. To expand the outlook of combination therapies, the objective of this study was to use high-throughput data generation technologies such as mass spectrometry and proteomics to investigate the response of HER2+ breast cancer cells to a mixture of two kinase inhibitors that has not been adopted yet as a standard treatment regime. The broader landscape of biological processes that are affected by inhibiting two major pathways that sustain the growth and survival of cancer cells, i.e., EGFR and PI3K/AKT, was investigated by treating SKBR3/HER2+ breast cancer cells with Lapatinib or a mixture of Lapatinib/Ipatasertib small molecule drugs. Changes in protein expression and/or activity in response to the drug treatments were assessed by using two complementary quantitative proteomic approaches based on peak area and peptide spectrum match measurements. Over 900 proteins matched by three unique peptide sequences (FDR<0.05) were affected by the exposure of cells to the drugs. The work corroborated the anti-proliferative activity of Lapatinib and Ipatasertib, and, in addition to cell cycle and growth arrest processes enabled the identification of several multi-functional proteins with roles in cancer-supportive hallmark processes. Among these, immune response, adhesion and migration emerged as particularly relevant to the ability to effectively suppress the proliferation and dissemination of cancer cells. The supplementation of Lapatinib with Ipatasertib further affected the expression or activity of additional transcription factors and proteins involved in gene expression, trafficking, DNA repair, and development of multidrug resistance. Furthermore, over fifty of the affected proteins represented approved or investigational targets in the DrugBank database, which through their protein-protein interaction networks can inform the selection of effective therapeutic partners. Altogether, our findings exposed a broad plethora of yet untapped opportunities that can be further explored for enhancing the anti-cancer effects of each drug as well as of many other multi-drug therapies that target the EGFR/ERBB2 and PI3K/AKT pathways. The data are available via ProteomeXchange with identifier PXD051094.

## Introduction

Breast cancer is a heterogeneous disease, and the HER2 positive subtype is characterized by the overexpression of the HER2 receptor in ∼20 % of breast cancers [1]. The HER2 receptor was among the first ones to be targeted, and trastuzumab was the first monoclonal antibody drug to be approved by FDA for HER2+ breast cancers. Despite the success achieved by trastuzumab treatment, alone or in combination with other drugs and chemotherapeutic agents, resistance to such therapies emerges within a year in most patients [2]. Many other therapeutic drugs have emerged since the discovery of trastuzumab in 1998 [3]. For example, Lapatinib and Ipatasertib (GDC-0068) are two drugs that have been developed to target the EGFR/HER2 receptors and pan-AKT kinases, respectively, and their downstream MAPK and PI3K/AKT signaling pathways [4,5]. Lapatinib is a reversible dual Tyr kinase inhibitor that targets both the HER2 and EGFR receptors by acting as an ATP-competitive small molecule that binds the intracellular catalytic region of the two kinases [4], while Ipatasertib, also a small molecule drug, inhibits the activity of all three Ser/Thr AKT kinase isomers [5,6]. Each of these drugs has shown promise in treating breast or other cancers alone or in combination with other therapeutic components [7–11]. In ∼50 % of HER2+ breast cancers the PI3K/AKT pathway is altered as well, its hyperactivation leading to alternative pathways that lead to the development of drug resistance [7]. Therefore, drugs that inhibit the kinases in this pathway are used in HER2+ breast cancers in combination with HER2 targeted therapies that rely on trastuzumab and Lapatinib [5–8]. In one study, combining Lapatinib and Ipatasertib has been shown to be beneficial to overcoming resistance to HER2+ therapy that evolved as a result of PIK3CA (Phosphoinositide 3-Kinase Catalytic Subunit Alpha) mutations [8]. The combination of Ipatasertib with Lapatinib has not been widely researched, however, and even less so through mass spectrometry (MS) technologies. Currently, NCI is listing 14 clinical trials that involve the use of Ipatasertib, in combination with either chemo- or immunotherapies for various cancer types [12]. The first pan-AKT inhibitor of hormone receptor positive/HER2+ negative advanced or metastatic breast cancers, Capivasertib, was approved only recently in 2023 by FDA, increasing therefore the interest toward AKT/PI3K targeted therapies.

The MAPK and PI3K/AKT pathways extensively cross-talk with each other through various mechanisms including cross-activation, cross-inhibition, and convergence [13], indicating the complexity of signaling networks that act within a cell. This presents challenges, but also untapped opportunities for the advancement of targeted therapies. Investigating the inhibition of MAPK and PI3K/AKT pathways can lead to an improved understanding of cancer progression and survival, as well as of the cross talk between the two pathways, which will be critical for the development of therapies that bypass the causes that lead to the development of drug resistance. This study was aimed at exploring the combinatorial effects of two small molecule inhibitors, Lapatinib and Ipatasertib, in HER2+ breast cancer cells, by using MS technologies. Mass spectrometry was utilized for its specificity, sensitivity, quantitative ability, and comprehensive profiling capability of the cellular proteome [14,15]. The response of SKBR3/HER2+ cells to Lapatinib and Ipatasertib was explored by proteomic profiling of cells exposed to 36 h inhibition with the drugs. Beyond the two targeted pathways, the results exposed a broad range of biological processes that were affected by the drug treatments, providing insights into the diverse landscape of proteins that can be explored for the development of future therapeutics and co-targeting strategies.

## Methods

### Reagents and materials

SKBR3 cells, representative of breast cancer cells overexpressing the HER2 receptor, were purchased from ATCC (Manassas, VA). Supporting reagents for cell culture, McCoy’s 5A (Modified), trypsin (0.25%)/EDTA (0.53 mM), Dulbecco’s Phosphate Buffered Saline (DPBS), PenStrep, and fetal bovine serum (FBS) were purchased from Gibco (Carlsbad, CA). Normocin was purchased from InvivoGen (San Diego, CA). Reagents used for sample processing such as NaF, Na_3_VO_4_, dithiothreitol (DTT), urea, ammonium bicarbonate, acetic acid, trifluoroacetic acid (TFA), protease inhibitor cocktail, Triton-X, Ribonuclease A (RNase A), and the nuclear/cytoplasmic cell fractionation Cell Lytic™ NuCLEAR™ extraction kit were from Sigma (St. Louis, MO). Sequencing grade trypsin was purchased from Promega (Madison, WI), and animal-Free Recombinant Human EGF from PeproTech (Cranbury, NJ). Propidium iodide was from Invitrogen (Waltham, MA). Lapatinib ditosylate and Ipatasertib were from Selleck (Houston, TX). For Western blotting, the antibodies were purchased from Cell Signaling Technology/Danvers/MA (Rabbit Primary mAbs CD82-D7G6H, VTCN1-D1M8I, PDCD4-D29C6, and HRP-conjugated Anti-Rabbit IgG Secondary Antibody) and R&D Systems/Minneapolis/MN (14-3-3-Sigma polyclonal goat IgG and HRP-conjugated Anti-Goat IgG Secondary Antibody). The buffer reagents were from Sigma (i.e., Trizma/TrizmaHCl glycine, sodium chloride, SDS, BSA fraction V, RIPA buffer), and the blot supplies from BioRad/Hercules/CA (Clarity Western ECL substrate, Mini-Protean TGX stain-free gels, Immun-Blot LF PVDF membrane and filter papers, Mini Trans-blot filter paper, Precision Plus protein standards/ladder, 2X Laemmli buffer, Tween 20, Blottin-grade blocker-nonfat dry milk). Zorbax SB300-C18/5 μm particles and SPEC-PTC18/SPEC-PTSCX cleanup pipette tips were purchased from Agilent Technologies (Santa Clara, CA). HPLC-grade solvents methanol and acetonitrile were purchased from Fisher Scientific (Fair Lawn, NJ). Cell culture materials including Nunc flasks and pipettes were from Life Technologies (Carlsbad, CA). Water was either produced by a MilliQ Ultrapure water system (Millipore, Bedford, MA) or distilled in-house from DI water.

### Cell culture and treatment with drugs

SKBR3 stock batches were generated from the original ATCC vial, and authentication was performed at ATCC by Short Tandem Repeat (STR) analysis. Normocin, a formulation of three antibiotics active against mycoplasma, bacteria and fungi, was added to the frozen cell stock. The SKBR3 cells were grown in T175 cm^2^ Nunc flasks in a water jacketed CO_2_ (5 %) incubator at 37 °C, in McCoy’s 5A medium supplemented with FBS (10 %). Upon reaching ∼70–80 % confluence, the cells were washed twice with DPBS (∼20 mL) and incubated in serum-free McCoy’s 5A for 24 h. Next, the cells were washed with McCoy’s 5A, and preconditioned for 15 min either with McCoy 5A serum-free medium (SFM) as control or with McCoy 5A supplemented with drugs (see concentrations below). Last, the cells used as control were incubated for 36 h with McCoy 5A in the presence of FBS (10 %) and EGF (10 nM), while cells exposed to the drugs were incubated with McCoy 5A/FBS (10 %)/EGF (10 nM) supplemented with Lapatinib (10 μM) alone or a combination of Lapatinib (10 μM) and Ipatasertib (1 μM). All cell cultures contained PenStrep (0.5 %). The cells were harvested by trypsinization (5-10 min), rinsed first with McCoy 5A/FBS (10 %) and then twice with cold PBS, pelleted at 500 x g (5 min, 4 °C), and frozen at -80 °C until further use. Three independent biological replicates were produced for each dug treatment (n=3).

### FACS analysis

FACS analysis of all control and drug-treated cells was performed to assess the cell cycle stage upon different drug regimens. Upon harvesting, a small portion of the cells were fixed in 70 % ethanol and preserved at -20 °C. Before analysis, the cells were washed once in DPBS and stained in a freshly prepared solution of 0.02 mg/mL propidium iodide, 0.2 mg/mL RNase A, and 0.1 % v/v Triton X-100 in DPBS. After 30 min incubation in the staining solution, in dark at room temperature, the samples were analyzed by a FACSCalibur flow cytometer (BD Biosciences, San Jose, CA).

### Cell fractionation and processing

To separate the cytoplasmic and nuclear cell fractions, the Cell Lytic™ NuCLEAR™ kit was used along with the protocols recommended by the manufacturer. Both cytoplasmic hypotonic lysis (HEPES 10 mM, pH 7.9, MgCl_2_ 1.5 mM, KCl 10 mM) and high-salt content nuclear extraction (HEPES 20 mM, pH 7.9, MgCl_2_ 1.5 mM, NaCl 0.42 M, EDTA 0.2 mM, glycerol 25 % v/v) buffer solutions were supplemented with phosphatase inhibitors (Na_3_VO_4_ and NaF, 1 mM each), a protease inhibitor cocktail (1 % of the extraction buffer), and DTT (1 mM). Briefly, the cells were initially allowed to swell with the hypotonic lysis buffer on ice for 15 min, and then vortexed for 10-15 sec with IGEPAL CA-630 (10 %, 0.6 % final concentration) for completing the lysis. The cytoplasmic extract was separated from the nuclear pellet by centrifugation at 10,000 x g for 1 min, and then the nuclei were disrupted with the high-salt content buffer by vortexing for 30 min. The nuclear extract supernatant was collected by centrifugation at 20,000 x g for 5 min. All operations were performed at 4 °C. The protein concentrations in each cell extract were measured with the Bradford assay (SmartSpec Plus spectrophotometer, Bio-Rad, Hercules, CA) using the Bradford dye reagent and bovine standards (BioRad, Hercules, CA). The cell extracts were further denatured/reduced with urea (8 M)/DTT (5 mM), pH∼8, for 1 h at 56 °C, diluted 10X with NH_4_HCO_3_ (50 mM), and digested with sequencing grade trypsin at protein:trypsin ratio of ∼50:1 w/w, pH∼7.8, overnight, at 37 °C. The enzymatic reaction was quenched the following day with TFA (1% of total cell extract volume). Prior to LC/MS analysis, the cell extract peptide mixtures were disposed of salts and detergents with SPEC-PTC18 and SPEC-SCX cartridges. The evaporated samples were reconstituted in a solution of H_2_O/CH_3_CN/TFA (98:2:0.01 v/v) and frozen at -80 °C until LC/MS analysis.

### Liquid chromatography (LC)-data-dependent analysis (DDA)-MS

The analysis of nuclear and cytoplasmic proteolytic digests was performed with an EASY nano-LC 1200 system and an EASY-Spray column ES902 (75 μm i.d. x 250 mm long, packed with 2 μm C18/silica particles, Thermo Fisher Scientific) that was operated at a flow rate of 250 nL/min at 45°C [16–19]. The mobile phases were prepared from H_2_O:CH_3_CN:TFA, mixed in proportions of 96:4:0.01 v/v for mobile phase A and 10:90:0.01 v/v for mobile phase B. A separation gradient of 125 min was used with the eluent B concentration increasing from 7 % to 30 % (105 min), 30-45 % (2 min), 45-60% (1 min), and 60-90 % (1 min), where it was kept for 10 min, and then decreased to a final concentration of 7 %. The mass spectrometry data were acquired with a QExactive/Orbitrap mass spectrometry system (Thermo Fisher Scientific) using a nano-electrospray ionization source (2.2 kV), a scan range of 400-1,600 m/z, resolution of 70,000, AGC target of 3E6, and maximum IT of 100 ms. For data-dependent MS2 acquisition (dd-MS2) with higher-energy collisional dissociation (HCD), the precursor ions were isolated with a width of 2.4 m/z and fragmented at 30 % normalized collision energy (NCE). MS2 resolution was set to 17,500, AGC target to 1E5 (minimum AGC target 2E3 and intensity threshold 4E4), maximum IT to 50 ms, and loop count to 20 [17–19]. Unassigned, 1+, and >6+ charges were excluded, apex trigger was set to 1 to 2 s, dynamic exclusion time to 10 s (peak widths of ∼8 s), and the isotope exclusion and preferred peptide match features were turned on. All samples were analyzed in triplicate (technical replicates), independent of each other, in random order within nuclear or cytoplasmic blocks from within each biological replicate.

### Targeted mass spectrometry

Parallel reaction monitoring (PRM) was utilized to validate the presence and change in abundance of selected peptides and proteins [17,18]. Peptide selection for PRM analysis was based on a lab-developed framework that assessed spectral quality based on XCorr scores, charge states, lack of PTMs, and the retention time (RT) relative standard deviations (SDs). The selected peptides were searched within a 20 min time-window of the peptide RT, following the same separation gradient as the original DDA-MS analysis. The precursor ions were isolated with a width of 2.0 m/z and fragmented at 30 % NCE with PRM parameters set as follows: resolution 35,000, AGC target 2E5, and maximum IT 110 ms. The PRM data were processed by the Skyline 20.2 software [20] by using a mass spectral library generated from the DDA-MS searches of the respective samples. The transition settings included the use of b and y ion types, from ion 1 to last ion, with precursor charges of 2, 3 and 4, and ion charges of 1, 2 and 3. The 5-10 most intense product ions were picked from the filtered product ions. The presence of a peptide was considered validated when the peptide displayed a minimum of 5 transitions and a dot product (dotp) score >0.9.

### MS data processing

The MS raw files were processed by the Proteome Discoverer 2.5 software package (Thermo Fisher Scientific, Waltham, MA) by using the Sequest HT search engine and a *Homo sapiens* database of 20,399 reviewed, non-redundant UniProt protein sequences (August 2022 download) [16–19]. The three LC-MS/MS technical replicates of each sample (control or treated cells, nuclear or cytoplasmic fractions) were combined in one multiconsensus protein and peptide report to increase the number of protein IDs and improve the quality of quantitative comparisons. The *Processing Workflow* search parameters included the followings: the Spectrum Selector peptide precursor mass range was set to 400-5,000 Da, the Sequest HT node parameters enabled the selection of 6-144 aa length fully tryptic peptides comprising maximum two missed cleavages, ion tolerances were set to 15 ppm for the precursor ion and 0.02 Da for the b/y/a ion fragments, and dynamic modifications were enabled for Met (15.995 Da/oxidation) and Nt (42.011 Da/acetyl) for all samples. The PSM Validator node used target/decoy concatenated databases with FDR targets of 0.01 (strict) and 0.03 (relaxed). Additional parameters were set in the *Consensus Workflow* for both peptide and protein levels. The PSM Filter eliminated all PSMs with Xcorr<1, the Peptide Group Modification site probability threshold was set to 75, and the Peptide Validator node to automatic with peptide level error rate control. All PSM, peptide, and protein FDRs were set to high (0.01) and medium (0.03). For Protein Grouping, the strict parsimony principle was enabled. Lastly, the Peptide/Protein filter node retained only peptides of at least medium confidence and proteins matched by only rank 1 peptides, and peptides were counted only for the top scoring proteins.

### Quantitation and statistical analysis

For assessing changes in protein expression, quantitation was performed by using either spectral counting or peak area measurements. Three biological replicates of drug-treated cells were compared to three biological replicates of EGF-control cells. The input for each biological replicate consisted of the multiconsensus report generated from the three technical replicates acquired for each sample. The reproducibility between any two sets of biological replicates was evaluated at the peptide and PSM levels based on RTs, XCorr scores, and total spectral counts.

### Spectral counting-based quantitation

PSM-based quantitation was performed based on the total PSM counts for a protein, with missing values being handled by adding one spectral count to each protein from the dataset. Normalization was performed at the global level by averaging the total spectral counts of the six samples taken into consideration (e.g., three drug-treated *vs* three EGF-treated samples) and using the resulting average as a correction factor for adjusting the counts of individual proteins in each sample. Differentially expressed proteins were selected by calculating the Log2 values of the spectral count ratios of the two datasets and using a two-tailed unpaired *t*-test for assessing significance. No data were excluded from the analysis, but only proteins matched by three unique peptides with fold change (FC)≥2 in spectral counts [i.e., Log2(Treatment/Control) either ≥1 or ≤(-1)] and p-value<0.05 were considered for discussion.

### Peak area-based quantitation

Area-based quantitation was performed with the aid of a Proteome Discoverer/*Label Free Quantitation (LFQ)* template that relied on a Percolator-based data *Processing Workflow* that used the same search parameters that were described above, with the exception of the PSM validator node that used the following settings: concatenated Target/Decoy selection, q-Value based validation, PSM maximum Delta Cn 0.05, and peptide Target FDRs of 0.01/0.05 (strict/relaxed). The Percolator-based workflow uses a semi-supervised learning algorithm and q-value based assessment of statistical significance to differentiate between correct and incorrect PSMs. The workflow also contained the Minora Feature Detector algorithm which detects and matches chromatographic peaks across LC/MS runs and links them to PSMs. The parameters for this node included: Minimum Trace Length 5, S/N Threshold 1, and PSM Confidence set to high. In the *LFQ* workflow, to account for peptide retention time shifts during multiple sample-runs on the LC column, chromatographic alignment was performed with the Feature Mapper node enabling a Maximum RT Shift of 15 min and maximum Mass Tolerance of 15 ppm. Normalization was performed based on total peptide abundance (all peptides used), and Protein Abundance calculations relied on the use of Precursor Ion Areas using the summed abundances of the connected peptide groups. Protein Ratio calculations were pairwise ratio based, i.e., using the median of all possible pairwise peptide ratios calculated between the replicates of all connected peptides. Modified peptides were excluded from the pairwise ratio-based quantifications. Low abundance resampling (lowest 5 % of detected values) was used as the mode of imputation for missing values. Differentially expressed proteins were selected based on the log2 values of the generated ratios (Treatment/Control) by applying a *t*-test. Proteins matched by three unique peptides with FC≥2 and abundance ratio p-value<0.05 were considered for discussion. Adjusted p-values accounting for multiple testing were calculated based on the Benjamini-Hochberg correction method.

### Bioinformatics data interpretation and visualization

A suite of software tools was used to place the results in biological perspective. Protein functionality was derived based on information provided by GeneCards [21] and UniProt [22]. STRING 11.5 was used to generate the networks of protein–protein interactions (PPI), and to assess enrichment in biological processes represented by the proteins that changed abundance [23]. The STRING interaction score confidences were set to medium and the process enrichment FDRs to <0.05. Cytoscape 3.8.2 and 3.9.1 software tools [24] were used to depict the PPI networks based on the interactomics report generated in STRING. Functionally related proteins were identified based on controlled vocabulary terms from UniProt. Cancer drug targets were extracted from the DrugBank database (Sep. 2021 download) [25]. RAWGraphs [26] was used for building the dendograms, bubblecharts, and the circular diagrams. Scatter plots of retention time and XCorr correlations were produced by Proteome Discoverer 2.5. All other figures were generated with Microsoft Excel.

### Western blotting

SDS-PAGE and Western blot experiments were performed by using a Mini-PROTEAN^®^ electrophoresis cell (Bio-Rad, Hercules, CA), precast stain-free gels, a Mini Trans-Blot® Cell Immun-Blot system, low fluorescence PVDF membrane/filter paper sets, enhanced HRP based chemiluminescence (ECL) protein detection on the blotting membrane, and a ChemiDoc^TM^ Imaging System (Bio-Rad). Whole cell lysates, up to 32 μg protein sample (20 μL) were loaded in each gel lane. All procedures followed the manufacturer’s instructions.

## Results

### Protein detection

The SKBR3 drug treatment strategy consisted of a multi-step process: (a) cell culture and synchronization in SFM for 24 h; (b) stimulation of control-cells with EGF (10 nM) for 36 h to assess canonical growth; (c) preconditioning of drug-treated cells for 15 min with drugs; and (d) treatment of cells with drugs for 36 h, i.e., Lapatinib (10 uM)/EGF (10 nM) or Lapatinib (10 uM)/Ipatasertib (1 uM)/EGF (10 nM) (**Figure 1A**). FBS (10 %) was added to all cell stimulations. The exposure of cells to drugs was performed for 36 h to allow for the observation of clear changes in the proteome profiles of drug-treated cells *vs* the EGF-control. After 36 h changes in cell morphology were observable, with some cells starting to detach. The proteome profiles of the harvested cells, separated into respective cytoplasmic and nuclear cell fractions, yielded ∼3,800-4,600 proteins per fraction, treatment, and replicate (**Figure 1Ba**). More than half of the detected proteins were identified by at least 2 peptides (**Figure 1Bb**) - group that was further considered for analysis - with protein identification reproducibility between three biological replicates being >75 % (**Figure 1Bc**).

**Figure 1:**
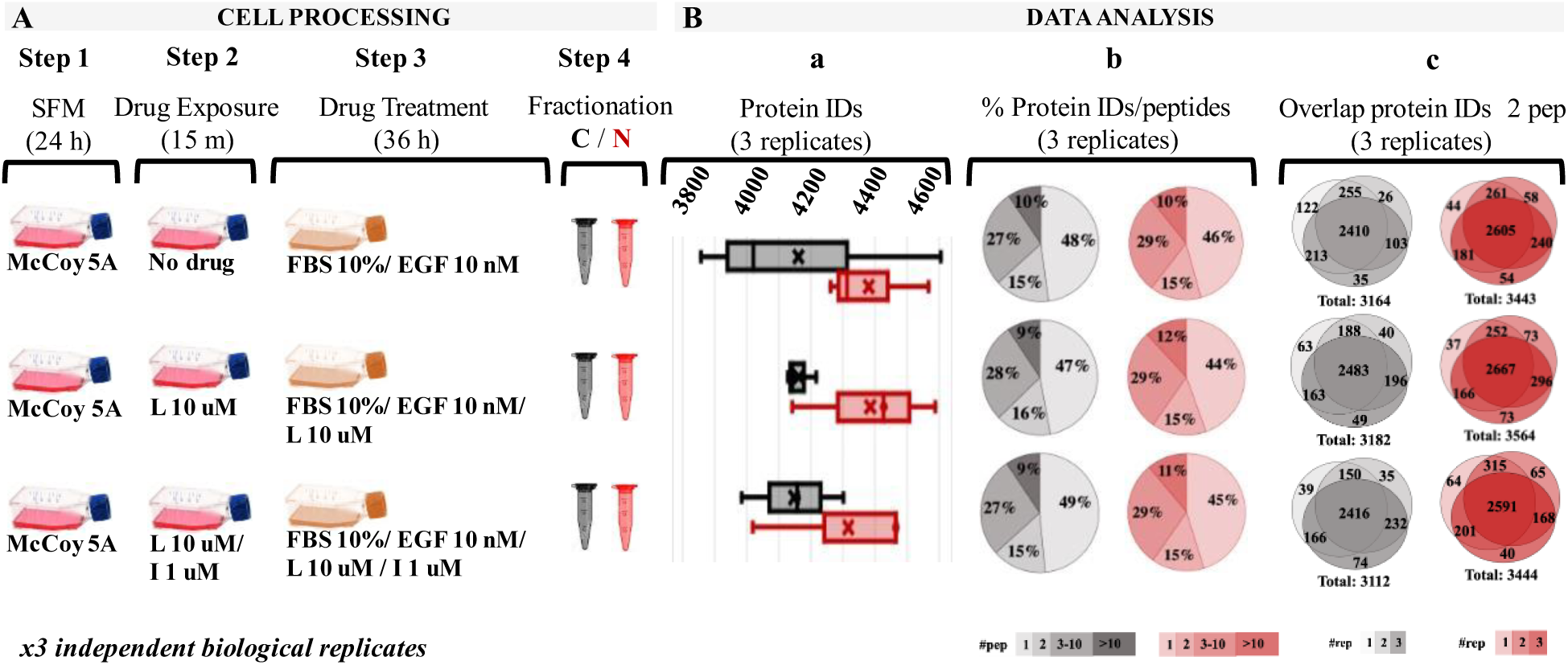
Overview of the drug treatment protocol and proteome profiling results based on spectral counting data. **(A)** Drug treatment steps; **(Ba)** Range of detected proteins in the three replicates of each cell fraction; (**Bb**) Pie charts representing the % distribution of detected proteins based on the number of unique peptides; (**Bc**) Protein identification reproducibility between three biological replicates (only proteins detected by at least 2 unique peptides were considered). Abbreviations: L - Lapatinib, I - Ipatasertib, C - cytoplasmic fraction (grey), N - nuclear fraction (red).

The reproducibility of peptide detection was assessed based on PSM counts, retention times, and XCorr scores. The PSM reproducibility for any two biological replicates of a specific treatment and fraction had a correlation score of 0.95 or higher (**Figures 2A-2F**, with results for the other measurements being provided in **Supplemental file 1**). Peptide XCorr and RT reproducibility across three different biological replicates, in the cytoplasmic and nuclear fractions of cells treated with EGF, as represented by the X-, Y-axes and color as the third dimension, is provided in **Figures 2G** and **2H**. The effectiveness of the nuclear/cytoplasmic separation was assessed based on the distribution of a set of nuclear (TOP1, TOP2A, TOP2B, LMNB1) and cytoplasmic (GAPDH, PKM, TUBB) protein markers in the two cell fractions (**Figures 2I** and **2J**). The nuclear markers were essentially localized to the nucleus (**Figure 2I**), while the cytoplasmic markers had a much higher abundance in the cytoplasmic than the nuclear cell fractions (**Figure 2J**). Their presence in the cell nucleus is supported by biological processes that sustain nucleocytoplasmic shuttling as well as the linkages that are established at the nucleo-/cytoskeletal interface [27]. The abundance of the nuclear/cytoplasmic markers was also representative of the broader range of ∼12,000 proteins identified in the whole dataset, with a distribution of the median PSM counts spanning over a range of ∼1-1,300 (**Figure 2K**). As expected, and as also confirmed by FACS, the 36 h exposure to drugs led the cells to a more pronounced G1 phase arrest when compared to the no-drug control (**Figure 2L**) (i.e., Control: 57-61 % G1, 24-26 % S, 4-11 % G2; Lapatinib treatment: 78-87 % G1, 7-16 % S, 2-10 % G1; Lapatinib/Ipatasertib treatment: 78-81 % G1, 6-11% S, 2-6 % G2). The PSM counts confirmed this result, the counts of the KI-67 nuclear proliferation marker dropping in the drug-treated cells to <30 % of their value in the EGF-control cells.

**Figure 2:**
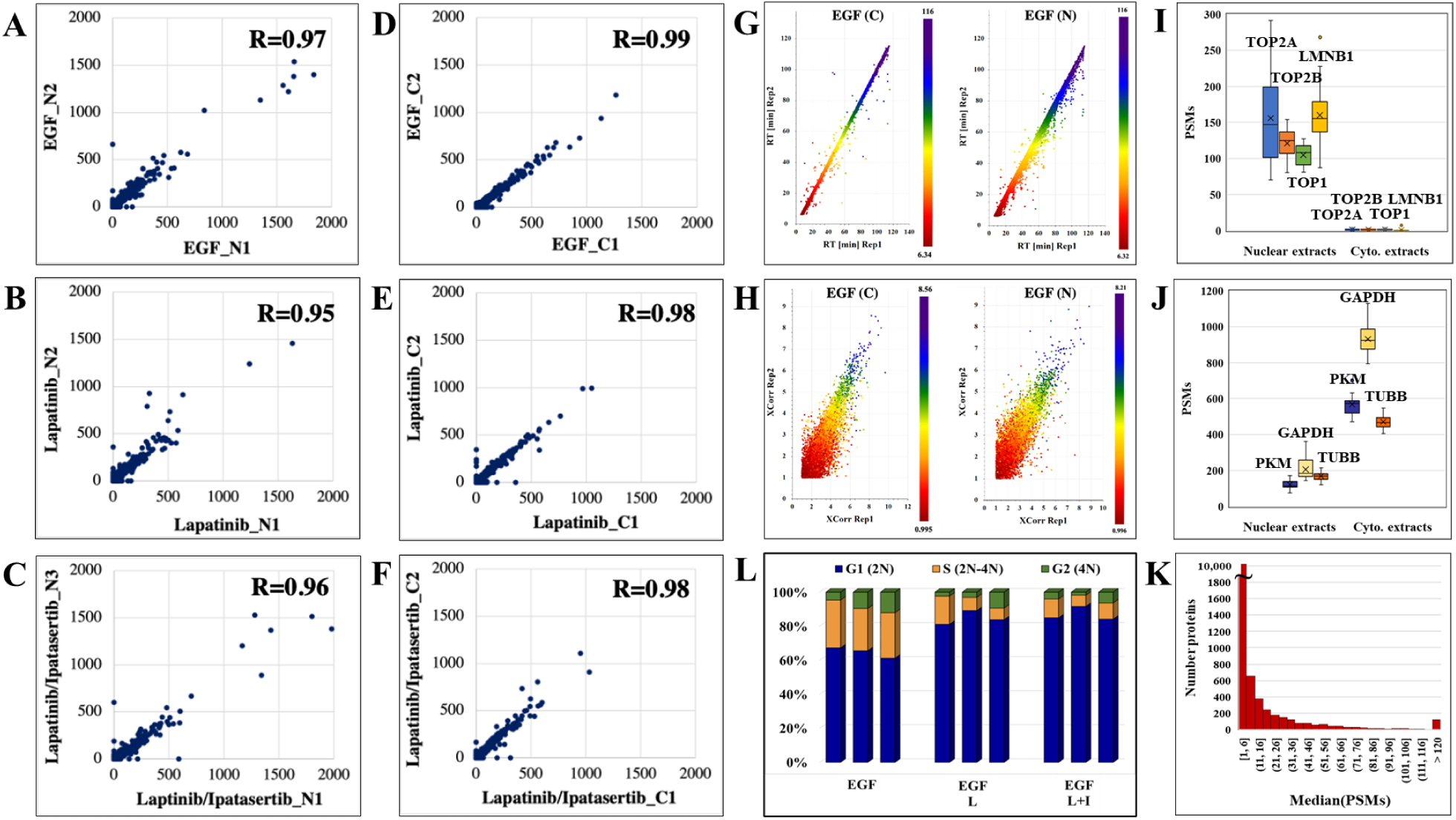
Data reproducibility. **(A-C)** Correlations between the peptide PSM counts in two biological replicates represented by the X-axis (replicate 1) and Y-axis (replicate 2 or 3) for the nuclear fractions of drug treatments. **(D-F)** Correlations between the peptide PSM counts in two biological replicates represented by the X-axis (replicate 1) and Y-axis (replicate 2 or 3) for the cytoplasmic fractions of drug treatments. (**G)** Correlations between the elution times of peptides detected in three biological replicates of the cytoplasmic and nuclear fractions of EGF-treated cells: replicate 1 (X-axis), replicate 2 (Y-axis), and replicate 3 (color bar). **(H)** Correlations between the XCorr scores of peptides detected in three biological replicates of the cytoplasmic and nuclear fractions of EGF-treated cells: replicate 1 (X-axis), replicate 2 (Y-axis), and replicate 3 (color bar). (**I-J**) PSM counts reflective of the abundance of nuclear and cytoplasmic markers in the nuclear and cytoplasmic cell fractions, respectively. (**K**) Histogram of the detected protein abundances in the whole dataset represented by the median of their PSM counts. **(L)** FACS distribution of cells after 36 h drug treatment in the G1, S, and G2 phases of the cell cycle. Abbreviations: L - Lapatinib, I - Ipatasertib, C - cytoplasmic fraction, N - nuclear fraction.

### Protein differential expression

The nuclear/cytoplasmic fractionation of cells resulted in the detection of two large protein groups, where each fraction was enriched by ∼70 % or more in the top 100 most abundant nuclear or cytoplasmic proteins, respectively, as calculated based on the number of matching peptides and as determined by the cellular compartment (CC) assignment in UniProt (**Figure 3A**). Two complementary label-free quantitative analysis approaches based on measuring the PSM counts and the peak areas were used to identify the group of differentially expressed proteins between treatment and control. The complementarity of these two methods has been previously documented, where the main advantage of the spectral counting method is the sampling of a larger range of protein abundances, while that of peak area measurements is the greater accuracy in assessing the protein abundance ratios with overlapping peptide ions [28].

**Figure 3.**
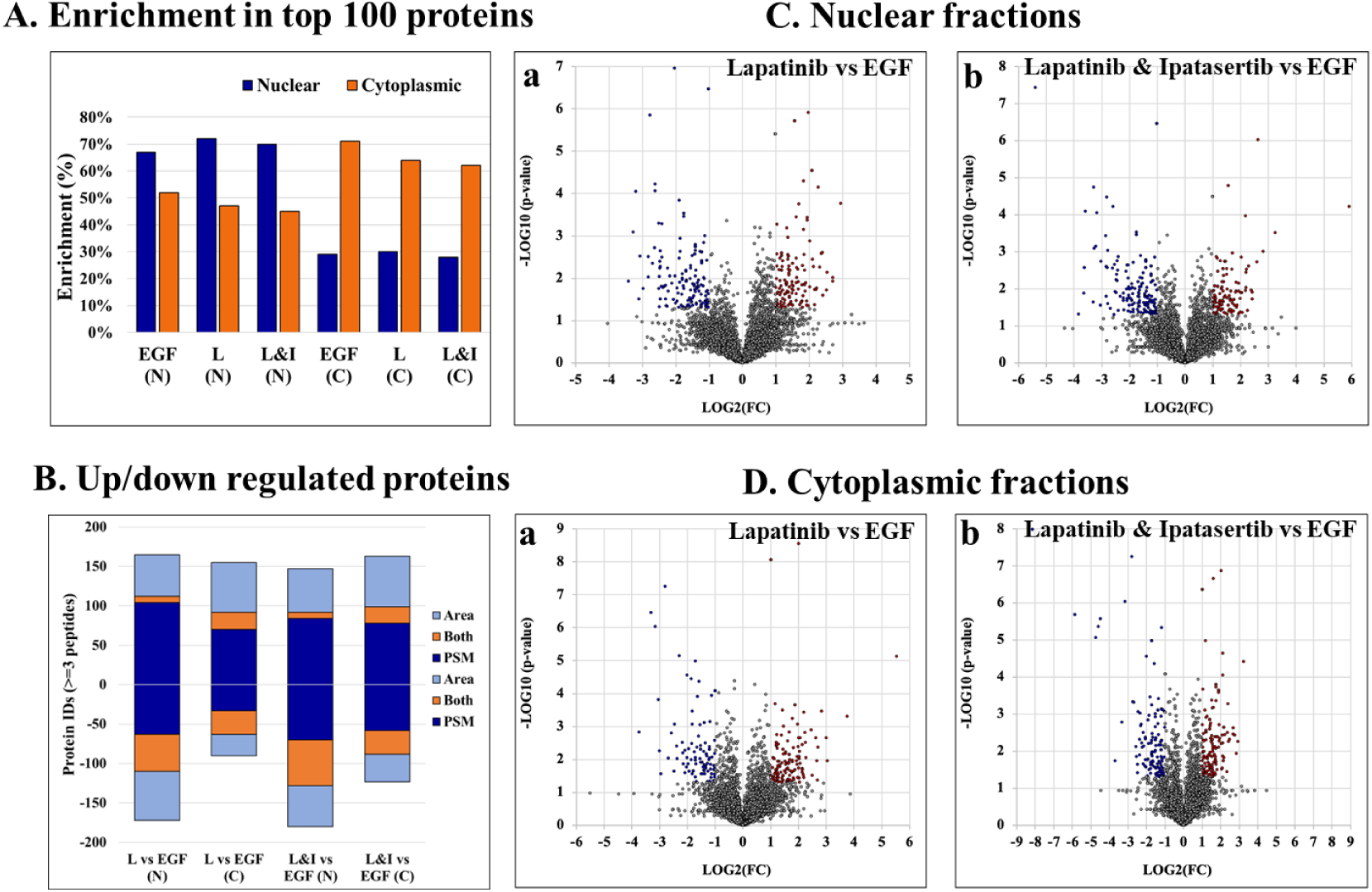
Results of the nuclear and cytoplasmic fraction enrichment process and differential expression analysis. **(A)** Nuclear and cytoplasmic protein enrichment among the top 100 most abundant proteins - as represented by total number of peptide matches per protein, calculated based on the CC location provided by UniProt. **(B)** Counts of differentially expressed proteins by each label-free quantification method. Up- and downregulated proteins are represented by positive or negative values, respectively. **(C)** and (**D**) Volcano plots representing differentially expressed proteins in the nuclear and cytoplasmic cell fractions, respectively, of Lapatinib (Ca, Da) and Lapatinib & Ipatasertib (Cb, Db) treatments compared to EGF-control. Differentially expressed proteins are indicated in red (upregulated) and blue (downregulated), and display ≥2-fold change abundance ratio with p-value ≤0.05. Abbreviations: L - Lapatinib, I - Ipatasertib, C - cytoplasmic fractions, N - nuclear fractions.

Proteins that were identified by at least 3 unique peptides and found to display at least 2-fold change in spectral counts or peak areas (FDR<0.05) were combined in a pool that was further evaluated in the study (**Figure 3B** and **Supplemental file 2**). Due to the higher sampling capacity of spectral counting, this method was utilized to construct the volcano plots and provide a better visual representation of the up- and downregulated proteins (**Figures 3C/3D** and **Supplemental file 1**). Overall, a total of ∼50-150 proteins per cell fraction displayed up-/or downregulation in response to the drug treatments (**Figure 3B**). A selection of biological processes that were represented by the proteins that changed abundance (i.e., ∼15 top processes comprising minimum 5 proteins per process, minimum fold enrichment of 5, and FDR<0.05), as captured by STRING and Gene Ontology (GO) analysis, is provided in **Figure 4** (full lists are provided in **Supplemental file 3)**. The drug treatments, either alone or in combination, resulted in the downregulation of cell cycle-related processes including G1/S and G2/M transition, spindle assembly and organization, chromosome segregation, and mitotic division, as largely represented by the nuclear proteins. These processes, along with other regulatory events of cell cycle progression, were also evident in the cytoplasmic fractions as represented by proteins with dual localization such as cyclin-dependent kinases (CDKs) or adapter proteins with broad implications in signaling such as 14-3-3 protein sigma. Other downregulated processes by the drug treatments included protein folding, adhesion, cytoskeletal/microtubule organization - as documented by the cytoplasmic proteins; and, translation, metabolism, and signaling in the nuclear and cytoplasmic fractions - as documented by kinases, adapters, and regulatory proteins. Lapatinib interrupts cell cycle progression and proliferative signaling by competitive inhibition of the ATP catalytic binding site of both EGFR and ERBB2 tyrosine kinases, while Ipatasertib induces cell cycle arrest and apoptosis by the inhibition of AKT Ser/Thr kinases (also known as protein kinase B or PKB). It has been shown to further complement this action by diminishing cell adhesion and invasive properties [10]. In breast cancers with amplified ERBB2 expression, hyperactivated AKT signaling was associated with the development of resistance to ERBB2 - targeted therapy [29]. Per Selleckchem manufacturer’s data, both drugs are highly selective, Lapatinib exhibiting greater than 300-fold selectivity for EGFR and ERBB2 over other kinases (i.e., c-Src, c-Raf, MEK, ERK, c-Fms, CDK1, CDK2, p38, Tie-2, and VEGFR2) and Ipatasertib displaying a 620-fold selectivity over PKA. In addition to inhibiting major events related to cell cycle progression and signal transduction, the treatment with drugs also resulted in the upregulation of processes that included chromosome and chromatin organization, DNA damage, splicing, and cellular senescence in the nuclear fractions, and broader cellular events related to metabolism, energy production, cellular respiration, transport and mitochondrial gene expression and organization in both nuclear and cytoplasmic fractions. When assessing the added effect of Ipatasertib, certain activities were found to be affected to a larger extent than in the Lapatinib treatment alone, including RTK, ERBB and MET signaling in the downregulated processes, and cellular senescence in the upregulated processes (**Supplemental file 3**). Ipatasertib further contributed to the downregulation of AKT targets and associated pathways such as mTOR and FoxO.

**Figure 4.**
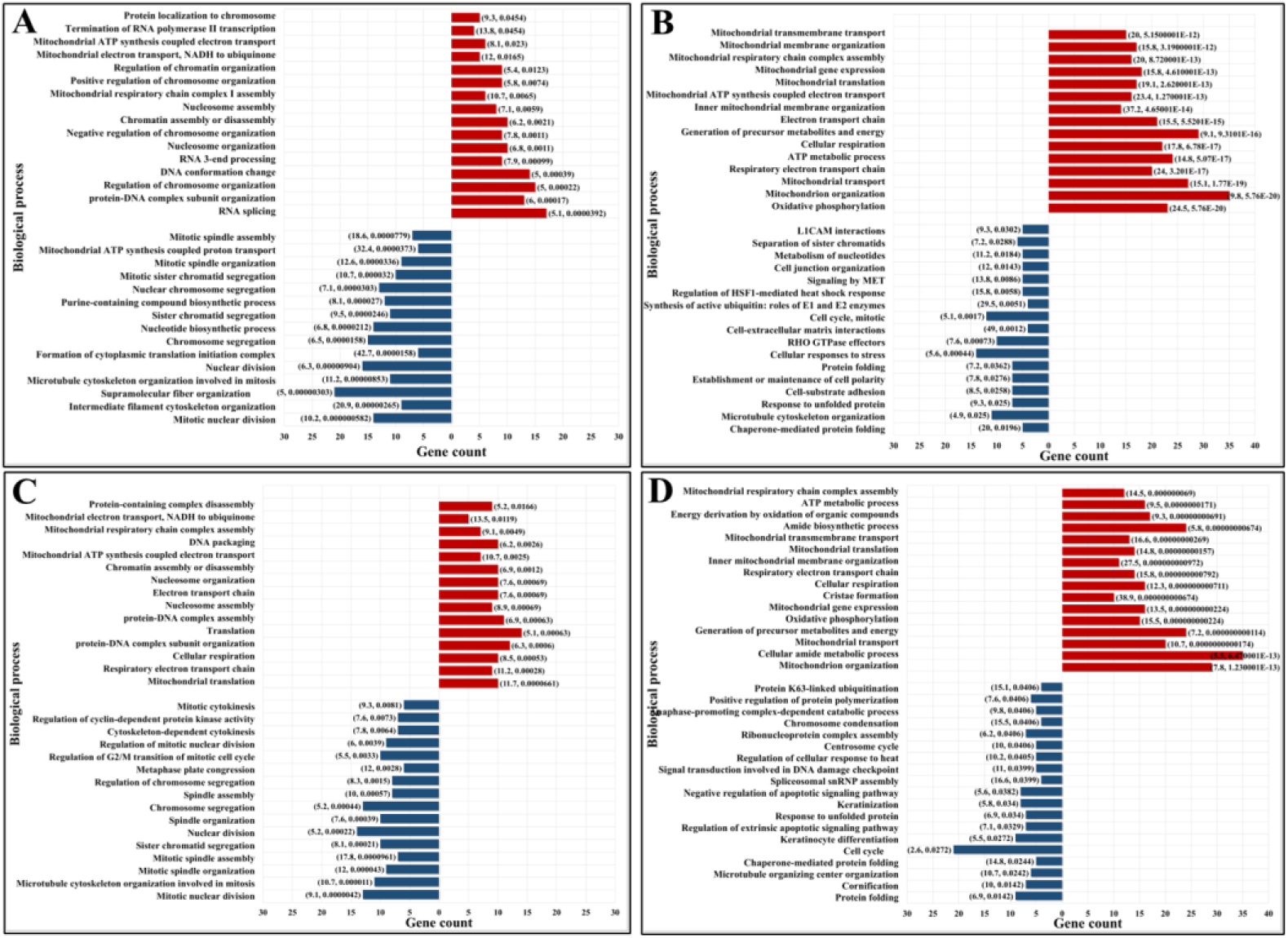
Enriched biological processes represented by the proteins that changed abundance in the drug-treated cells. **(A)** Lapatinib treated cells-nuclear fraction; (**B**) Lapatinib treated cells-cytoplasmic fraction; **(C)** Lapatinib/Ipatasertib treated cells-nuclear fraction; (**D**) Lapatinib/Ipatasertib treated cells-cytoplasmic fraction. Notes. The selected biological processes display at least 5-fold enrichment (FDR ≤0.05) and are represented by at least 5 genes/category. Upregulated processes are shown in red and downregulated processes in blue. The numbers in parenthesis indicate fold-enrichment followed by FDR.

To inform about the broader implications of the drug treatments, the protein-protein interaction (PPI) networks of the proteins that changed abundance in the nuclear fractions, which displayed a greater variety of up- and downregulated processes, are shown in **Figures 5** and **6**. On a background of cytoskeletal protein interactions, the PPI networks with the most interconnected nodes were tied to the above described gene expression, cell cycle and adhesion processes. Downregulated cellular respiration and metabolic processes formed additional clusters.

**Figure 5.**
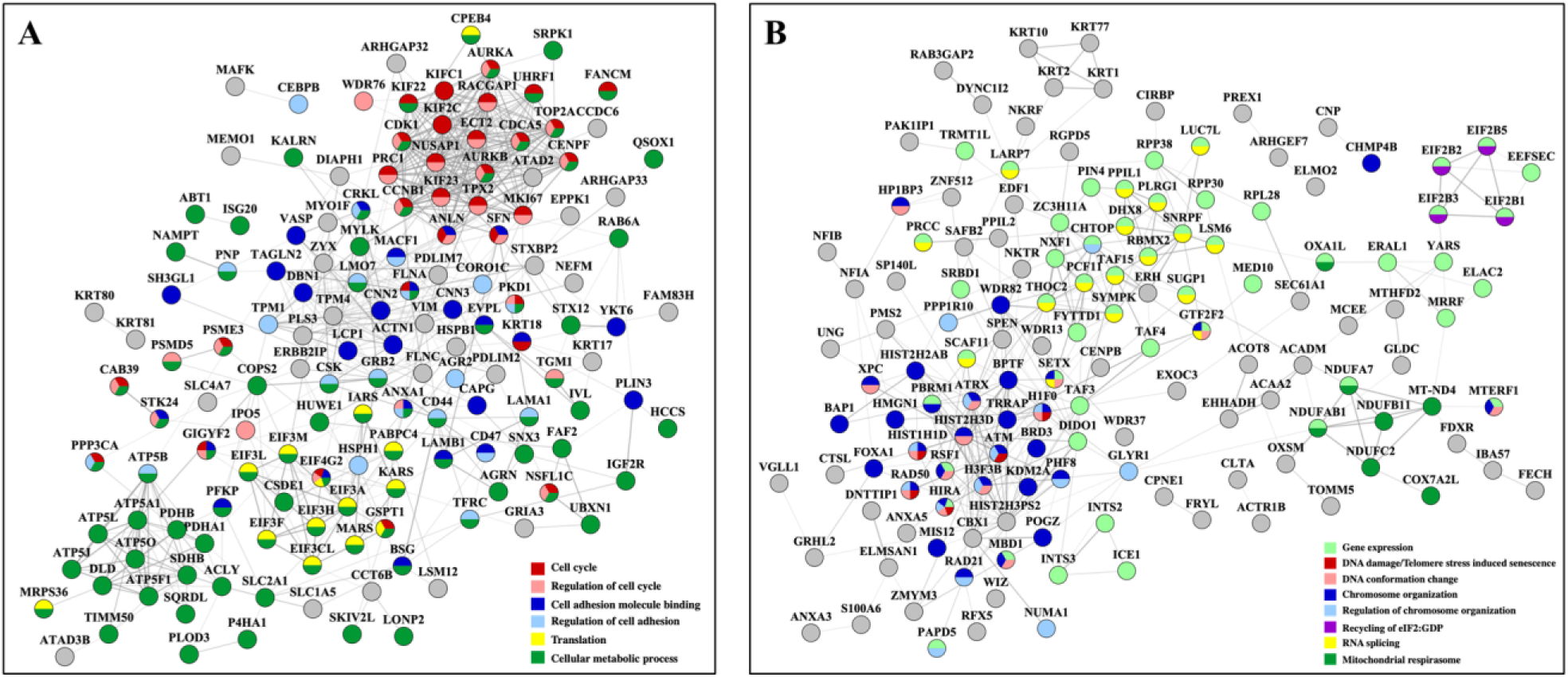
PPI networks of nuclear proteins that changed abundance in the Lapatinib-treated cells. (**A**) Downregulated proteins; (**B**) Upregulated proteins. Notes. The PPI networks were created with STRING and visualized in Cytoscape 3.9.0.

**Figure 6.**
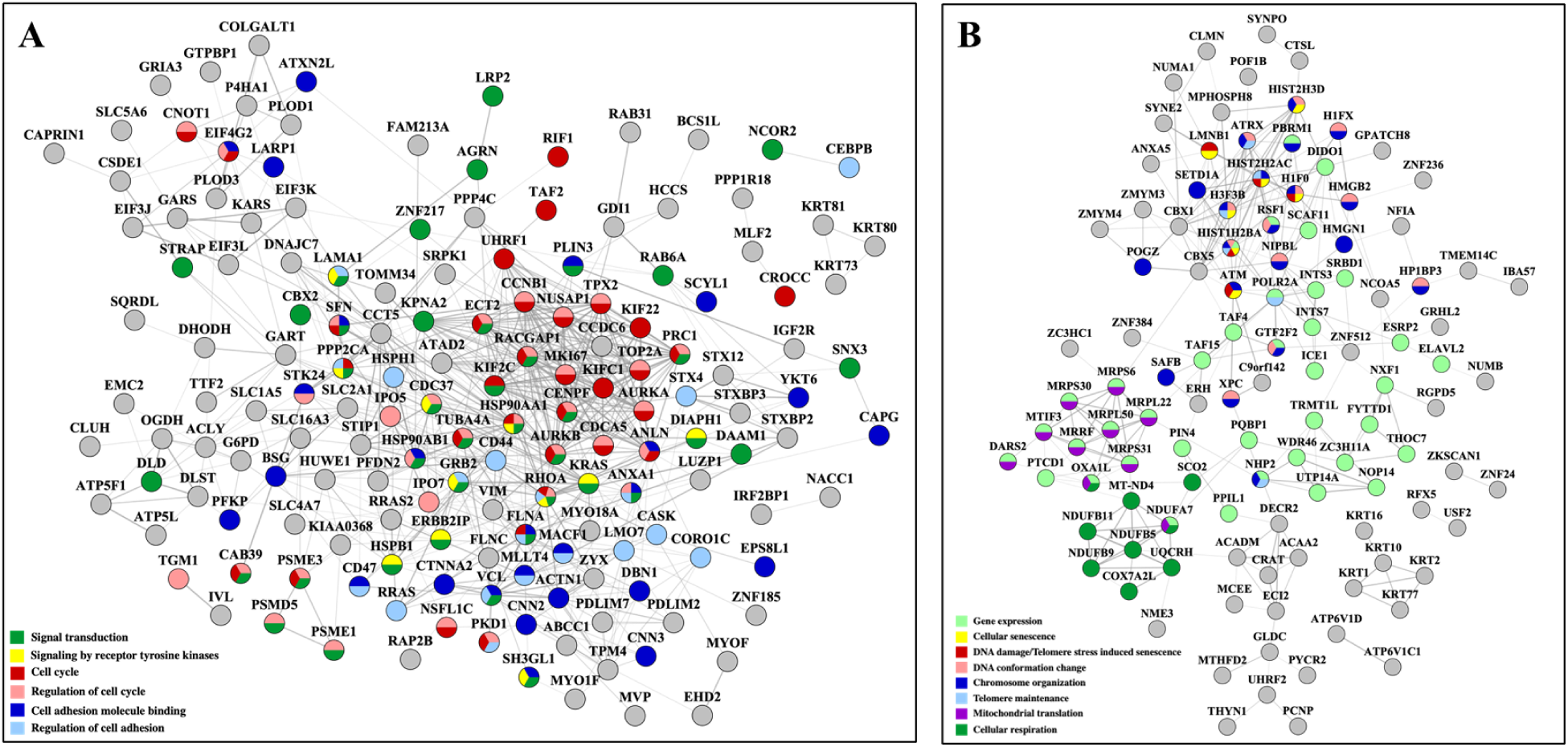
PPI networks of nuclear proteins that changed abundance in the Lapatinib/Ipatasertib-treated cells. (**A**) Downregulated proteins; (**B**) Upregulated proteins. Notes. The PPI networks were created with STRING and visualized in Cytoscape 3.9.0.

As some of the above-described biological processes were also representative of the cancer hallmarks, the differentially expressed proteins (altogether up/downregulated in either cytoplasmic or nuclear fractions and in either treatments) were matched to the ten hallmarks by using a previously in-house developed database of hallmark-related proteins [19] (**Figure 7A**). In agreement with the processes described above, the hallmarks that were mostly affected included cell communication/signaling, cell cycle/proliferation, cell death/apoptosis (with many overlapping proteins matching the proliferation category), and adhesion/motility (**Supplemental file 4**). Three additional categories of promise to advancing the understanding of the mechanism of action of these two drugs emerged, and included proteins with transcription factor activity, proteins involved in DNA damage response, and proteins involved in modulating inflammatory and immune systems processes. Moreover, a fairly large number of the affected proteins were found to have been previously catalogued in the Human Cancer Metastasis Database (HCMDB) [30] and the expert-curated COSMIC Cancer Gene Census (CGC) database of mutated genes that drive human cancer [31], underscoring the relevance of their role in cancer progression. Specific proteins with altered behavior that were part of the EGFR/ERBB2 and PI3K/PKB pathways that were targeted by the Lapatinib and Ipatasetib kinase inhibitors, along with the associated hallmark processes, are depicted in the circular diagram from **Figure 7B** and **Supplemental file 4**.

**Figure 7.**
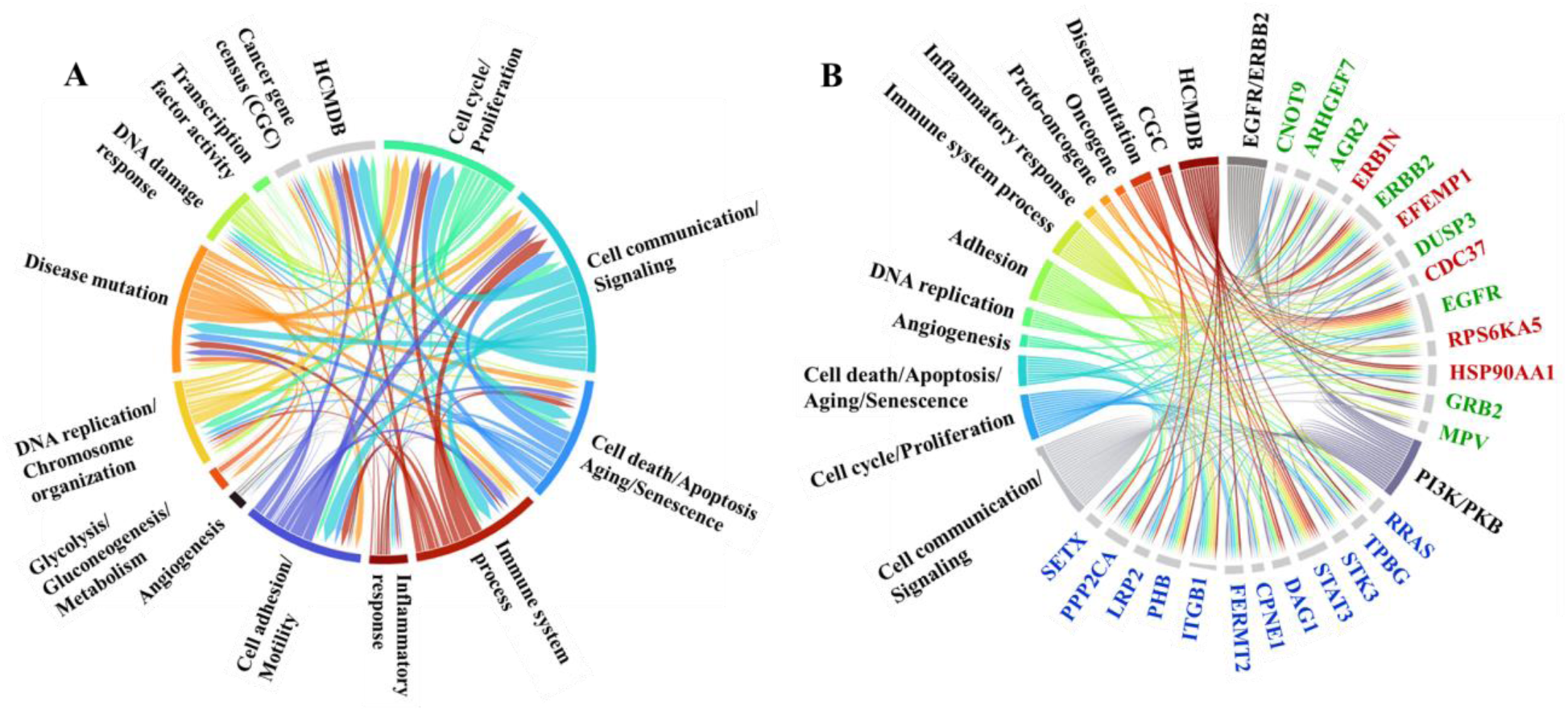
Circular diagrams illustrating the cancer-supportive biological processes that were affected by the drug treatments. (**A**) Cancer hallmarks, HCMDB and CGC database protein matches. (**B**) Proteins with altered behavior mapped to the EGFR/ERBB2 and PI3K/PKB pathways and associated cancer hallmarks. Gene name color code: green-proteins mapped to both pathways; red-protein mapped to the EGFR/ERBB2 pathway only; blue-proteins mapped to the PI3K/PKB pathway only.

Differentially expressed proteins for which consistent results [i.e., 3 unique peptides, log2(FC)>=1 or <=(-1), p<0.05] were obtained by at least three out of the four measurements in either the cytoplasmic or nuclear fractions (i.e., PSM and area-based measurements in both Lapatinib and Ipatasertib treatments) are represented in the bubble charts from **Figure 8A/B** and **Supplemental file 5.** Differences in fold-change abundances induced by the cell treatments are inferable from the figure [note bullet size for each protein and treatment]. A small protein subset that met the differential expression filtering criteria by both area and PSM measurements only in the Lapatinib/Ipatasertib cell treatments, but not Lapatinib alone, is shown in **Figure 9A**. This subset included the MRP1 protein (Multidrug resistance-associated protein 1) which is an ATP-dependent transporter that confers resistance to anticancer drugs (ABCC1). In addition, proteins with relevance to aberrant proliferation (oncogenes and tumor suppressors), metastasis (EMT and angiogenesis regulators), and diagnostics (epithelial and mesenchymal markers), and that emerged from either the Lapatinib or Lapatinib/Ipatasertib treatments are highlighted in **Figure 9B**. Their change in abundance or activity was supported by two or more measurements, the average of which being depicted in the figure. Several proteins representative of the above processes were validated by independent methods such as parallel-reaction monitoring (PRM)/MS or Western blotting (**Supplemental files 6** and **7**).

**Figure 8.**
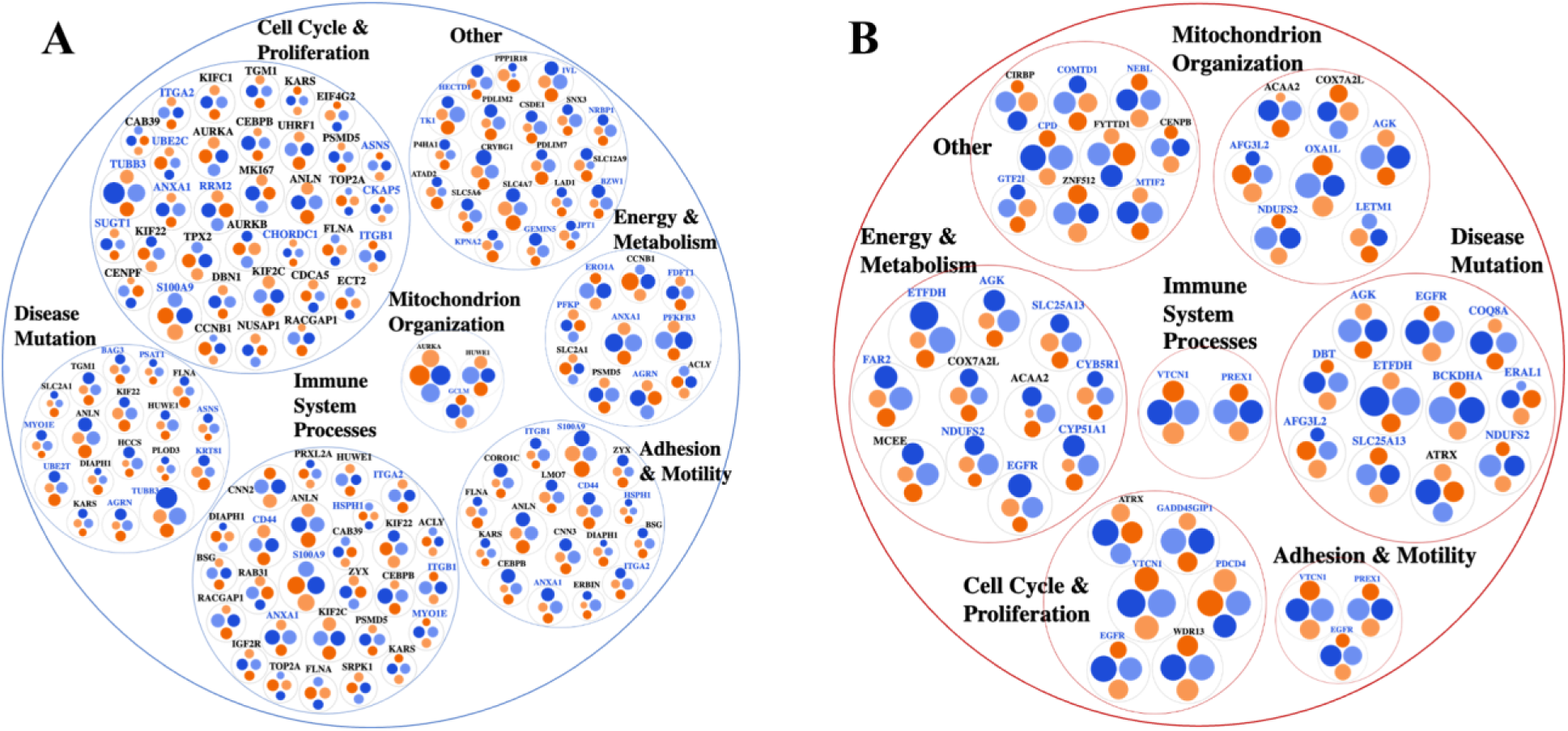
Bubble charts representing biological processes descriptive of cancer hallmarks that encompassed larger subsets of proteins with change in expression or activity in the nuclear (black font) or cytoplasmic (blue font) cell fractions (supported by at least three out of the four area or PSM comparisons). (**A**) Proteins with decreased counts. (**B**) Proteins with increased counts. Bullet color code: Lapatinib *vs* EGF measured by PSMs (light blue) and Area (light orange), and Lapatinib&Ipatasertib *vs* EGF measured by PSMs (dark blue) and Area (dark orange). The size of the node is proportional to the log2(FC) in each comparison.

**Figure 9.**
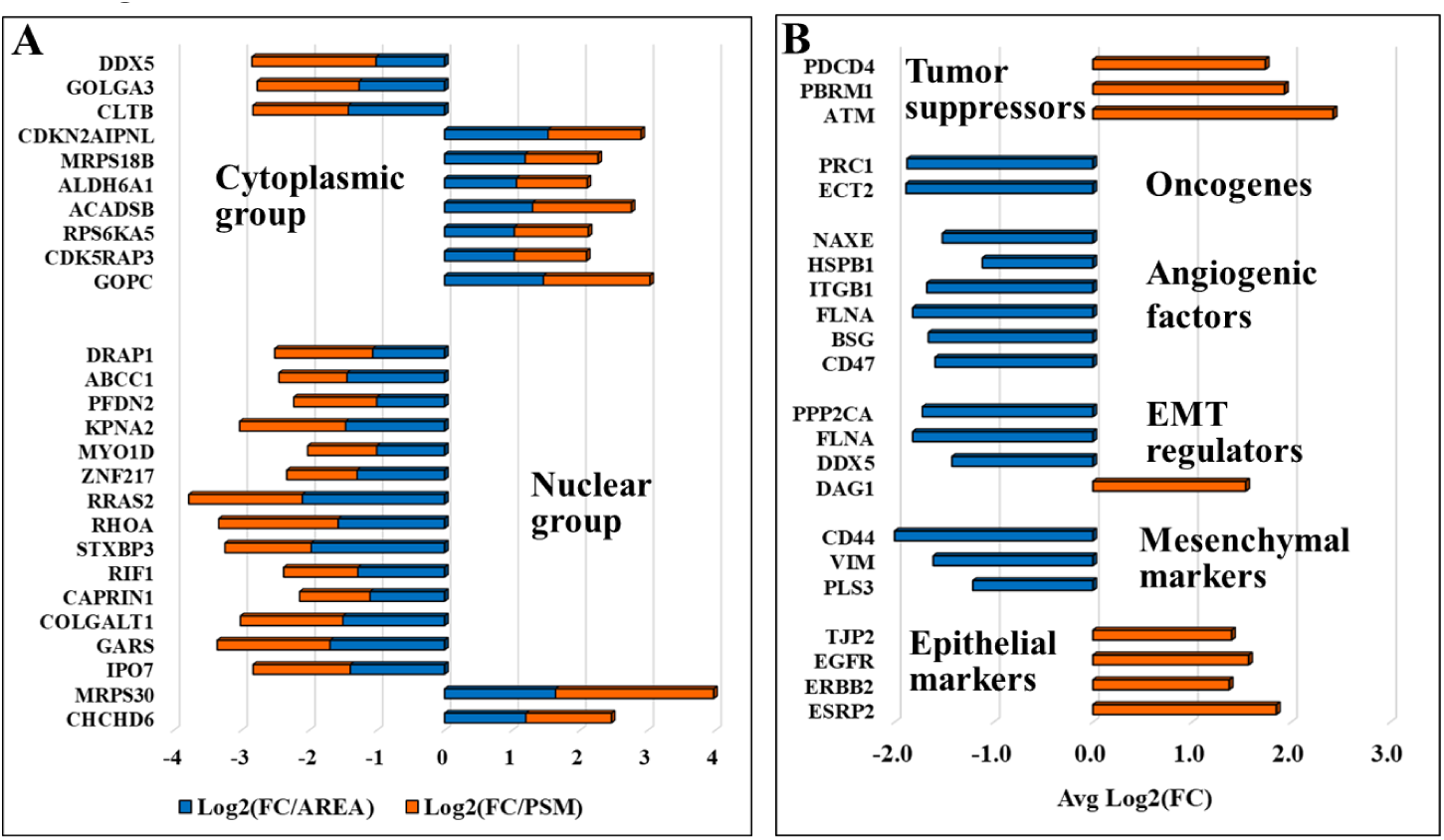
(**A**) Protein set that exhibited a change in abundance or activity in the Lapatinib/Ipatasertib cell treatments, but not in the Lapatinib treatments. (**B**) Selected proteins with relevance to aberrant proliferation, metastasis, and diagnostics.

Last, the differentially expressed proteins were further screened for their cancer drug targeting potential, based on information extracted from the DrugBank database of approved and investigational targets **Figure 10**) [25]. The analysis revealed that these proteins represented not only currently approved drug targets (28) but also investigational targets (43), still undergoing research on their potential therapeutic utility and effectiveness in the context of combination therapies that present interest for targeting alternative biological pathways at lower effective dosages (e.g., AURKA, AURKB, CDK1, CDK2, TOP2A) and for counteracting the proliferation of drug resistant cancer cells.

**Figure 10.**
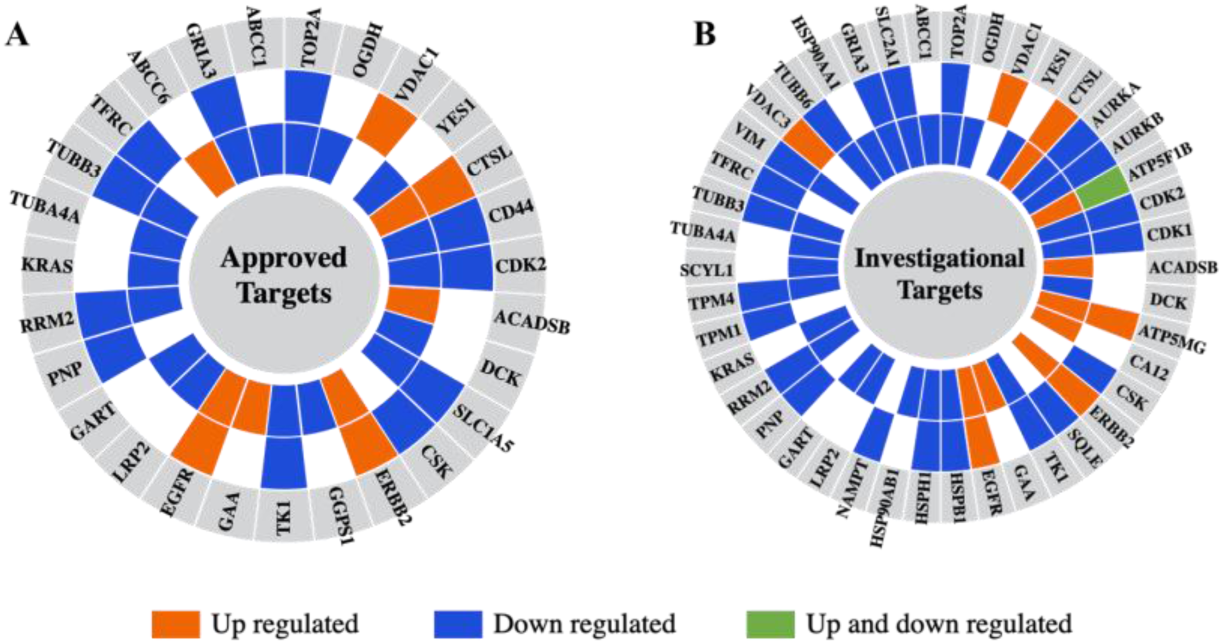
Differentially expressed proteins in the nuclear and cytoplasmic cell fractions aligned with approved **(A)** and investigational **(B)** drug targets from the DrugBank database. From the outer to the inner circle the comparisons are Lapatinib *vs* EGF and Lapatinib/Ipatasertib *vs* EGF. The different colors represent proteins that are upregulated (orange) or downregulated (blue) in either the nuclear or cytoplasmic cell fractions, or upregulated in one fraction and downregulated in the other (green).

## Discussion

The proteomic analysis of SKBR3/HER2+ breast cancer cells treated with the EGFR/ERBB2 receptor Tyr kinase inhibitor Lapatinib and pan-AKT Ser/Thr kinase inhibitor Ipatasertib revealed that the drugs exerted a wide-ranging impact on signaling and biological processes that support cancer progression (**Figures 4-9, Supplemental file 3**). The effect on the target EGFR/ERBB2 and AKT/PKB pathways was confirmed by the affected proteins that are part of these pathways and that further implicated the whole range of cancer enabling hallmarks (**Figure 7**). Broadly, the two treatments affected several signaling pathways leading to the downregulation of cell cycle progression, growth and division processes (MAPK, ERK1/ERK2, JAK/STAT), of pathways involved in immune responses (NF-kappaB, TNF-alpha), and of mechanisms supportive of adhesion, migration, and metastasis. Chromosome organization, transcription regulation, and DNA damage repair were among the upregulated processes. It should be emphasized that changes in PTM status or subcellular localization are common for many proteins involved in biological signaling and gene expression processes creating the artifactual impression of changes in expression level. Likewise, cell-membrane proteins often present altered glycosylation profiles and structural features that affect their consistent detectability [32]. Based on the results of the measurements, we will continue to refer to the selected proteins as being up- or downregulated, however, it must be understood that while the measurements reflect a drug-induced effect, this effect may represent a more complex and refined consequence that goes beyond a simple change in expression.

### Cell cycle, growth, proliferation

Not surprisingly, in the nucleus, Lapatinib treatment led to the downregulation of the cell proliferation and cancer prognosis maker (KI-67), G2/mitotic-specific cyclin-B1 (CCNB1), and cell cycle G1/S and G2/M transition as a whole as represented by a highly interconnected network of nuclear proteins (AURKA, AURKB, CDK1, CDCA5, CCNB1, TOP2A, SFN, TPX2, KIF2C, KIFC1, KIF22, KIF23, NUSAP1, PRC1, ECT2, RACGAP1, FANCM, ANLN, UHRF1) (**Figures 5**, **6, 8**). Many of these proteins are overexpressed in various cancer types and are linked to poor patient prognosis, tumor growth, invasion, and metastasis through signaling pathways such as MAPK, PI3K/AKT, and AMPK [33,34]. Proteins such as the mitotic Ser/Thr kinase cell cycle progression regulators AURKA and AURKB, cyclin CCNB1, and kinesin-like KIF2 displayed some of the largest and consistent changes in abundance. Anillin (ANLN) was the protein that displayed the largest downregulation in both the Lapatinib and Lapatinib/Ipatasertib drug treatments. ANLN is an actin-binding protein that is required in cytokinesis, localized at the cleavage furrow with roles in preserving its structural integrity, that was suggested as a potential target for cancer treatment [35]. 14-3-3 sigma or stratifin (SFN), found to be dually located in this work, has shown aberrant expression in various cancers, being often downregulated, but its upregulation has been associated with the development of resistance to therapeutic drugs and ECM remodeling [36]. SFN is an adapter protein that modulates the function of a broad range of proteins via binding, having thus regulatory roles in many signaling pathways and biological processes such as cell cycle, apoptosis, metabolism, protein trafficking, cell adhesion and motility [36,37]. Interestingly, 14-3-3 was found to protect against tumorigenesis by negatively regulating the activity of PKB [36]. Cytoplasmic proteins with regulatory roles in cell cycle progression were also downregulated by Lapatinib, including the proliferation marker TK1, PFKFB3 (regulator of CDK1 detected in the nuclear fraction), SFN, UBE2C, and SUGT1 required for G1/S and G2/M transition. An interesting finding was the downregulation of involucrin (IVL), an early differentiation marker of keratinocytes regulated by EGFR, which in contrast to our study was reported to be upregulated by Lapatinib [38]. The downregulation of proteins with high fold-change such as TK1 and PFKFB3 agreed with previous studies [39,40]. The largest change in expression in the cytoplasmic fraction was observed for TUBB3 (tubulin beta III) - a microtubule protein with important roles in chromosome segregation during mitosis, which was shown to be overexpressed in some cancers in a cell-cycle dependent manner, its overexpression leading to resistance to taxane derivates [41], advanced tumor stage, and HER2 amplification in patient cohorts [42]. The mechanisms that lead to TUBB3 expression dysregulation in cancer cells have not been, however, clarified yet [43].

### Immune response and inflammation

The impact of drug treatments on inflammatory and immune response processes was underscored by proteins that displayed downregulation in both subcellular fractions (e.g., S100A9, TBK1, ANXA1, KIF2C, CEBPB) (**Figures 7, 8**). S100A9 in the cytoplasmic fraction, and KIF2C in the nuclear fraction, displayed the largest change in abundance, both being also implicated in cell cycle progression. Overexpression of S100A9 has been implicated in the development and progression of many cancer types, including breast cancer, and correlated with high expression of Ki67 and HER2 [44]. The involvement of S100A9 in multiple cellular processes such as invasion (through H-Ras pathway), apoptosis (through p53 dependent or independent pathways), or inflammation (through NFKB pathway) has made S100A9 a protein of interest to many studies, which led ultimately to conflicting reports about its role in tumorigenesis [44,45]. MAPK and JAK1/2 inhibitors, and STAT3 silencing, have led to S100A9 expression inhibition [44,46] but to our knowledge no direct link was yet established between Lapatinib and downregulation of S100A9 expression. The overexpression of KIF2C in many cancers has led to proliferation, EMT activation, invasion, and metastasis via PI3K/AKT, mTORC1, and MAPK/ERK signaling [47–49]. Due to its multifunctional role in cell cycle related processes and genomic stability, it was reported that the suppression of KIF2C inhibits mitosis [47]. Yet, an important and less studied aspect is the involvement of KIF2C in immune responses, where the role of KIF2C in immune cell infiltration leading to tumor inhibition is either positively or negatively associated, depending on the cancer type [48,49]. Liu *et al.* reported that in breast cancer increased expression of KIF2C is associated with higher immune cell infiltration [49]. As such, investigating the behavior of infiltrating immune cells under Lapatinib treatment or Lapatinib/immunotherapy combinations could lead to new and more effective therapeutic applications of Lapatinib. Lastly, both S100A9 and KIF2C, as being an integral part of the tumor microenvironment, have been reported as suitable markers for evaluating immunotherapy response [49–51].

In the nuclear fractions, the altered behavior of CEBPB, a transcription factor involved in cell proliferation and regulation of inflammatory and immune responses [21], further underscored the broad impact exerted by the drug treatments. Previous studies have shown that the expression of CEBPB is not altered much in breast cancer *vs* normal cells. Nonetheless, changes in mRNA expression were observed between different cancer types, with increased levels being associated with metastatic and high tumor grade breast cancers [52]. To note, however, that the activity and subcellular localization of CEBPB is regulated by posttranslational modifications (phosphorylation, acetylation, methylation, sumoylation) that have been shown to alter the expression of various CEBPB isoforms [52]. The drug-induced inhibition of EGFR signaling in this study may have affected, therefore, the activity of this protein rather than the actual expression level [52]. Interestingly, in the cytoplasmic fraction, Annexin 1 (ANXA1) with known anti-inflammatory activity and loss of function or expression in cancer cells, was also downregulated [21]. The protein is implicated however in a broad range of processes that include chemotaxis, proliferation, cell adhesion, and motility, rendering the interpretation of drug treatment results more difficult [21].

### Adhesion and migration signaling

Along with processes that remodel the cytoskeleton, biological adhesion as reflected by cell-ECM and cell-cell interactions, focal adhesion, cell junction organization, anchoring junction, CAM and cadherin binding, and L1CAM interaction processes, was affected by a large number of proteins that displayed downregulation (e.g., CD44, ITGB1, ITGA2, TUBB3, TUBB6, PXN, ANXA1, DIAPH2, MARK2, LRRC15, BAG3, BZW1, CNN2, CNN3, SH3GL1, CD44, PLIN3, LAMA1, FLNA, LMO7, DBN1, ANXA1) (**Figures 4-8**). As cell-adhesion molecules can shuttle between transmembrane regions or from the cytosol to the nucleus [53], the change in abundance for several proteins was detected in the nuclear cell fractions. Some of the proteins with the highest fold-change such as CD44, calponins CNN2/3, and endophilin SH3GL1 exert multiple roles within a cell. CD44 is a multifunctional cell-membrane adhesion receptor that promotes tumor progression, invasion, metastasis, and epithelial-mesenchymal transition [54] through various mechanisms including nuclear translocation [55]. It is a stemness marker used for predicting metastatic propensity and also a therapeutic target [56]. Furthermore, CD44 has been demonstrated to play a role in the development of drug resistance in breast cancer cells treated with Trastuzumab, Lapatinib, and Tamoxifen [57–59], therefore targeting CD44 in conjunction with other potential drug targets remains of high interest. The homologous genes CNN2 and CNN3 are actin binding proteins with functions in cytoskeleton stabilization and processes related to cell proliferation, adhesion, migration, and metastasis [60]. These proteins are overexpressed in many cancer types but reports about their function in the cell are confounding depending on the cancer type [61–66]. Most studies state, however, that CNN2/3 overexpression promotes tumorigenesis and invasion probably through MAPK/ERK signaling [64]. In this work, these proteins were downregulated by the Lapatinib treatment. The protein levels of SH3GL1 (SH3 Domain Containing GRB2 Like 1, Endophilin A2) were linked to tumor progression, metastasis, and worse patient outcomes in many cancer types [67,68]. Endophilin silencing in SKBR3 cells led to impaired HER2 receptor internalization and reduced downstream signaling in cells treated with Trastuzumab, and reduced cytotoxicity in HER2+ cells treated with TDM-1 [68]. To our knowledge, the downregulation of these proteins by Lapatinib in SKBR3 cells is firstly reported here, providing novel opportunities for future research into the mechanism of action of this drug and its effects on biological processes such as adhesion and migration. Similar to the calponins, Coronin-1C (CORO1C), another actin filament binding protein with roles in cell cycle progression, apoptosis and cell adhesion/migration, has been shown to promote metastasis, its overexpression being associated with poor prognosis [69–71].

### Upregulated biological processes

Several nuclear proteins involved in chromosome organization, chromatin remodeling, transcription regulation, and DNA damage sensing and repair displayed altered upregulation or activation in response to the drugs (note the highly interacting cluster of proteins in **Figure 5B**: ATM, ATRX, XPS, FOXA1, MIS12, H1F0, RSF1, NUMA1, TRRAP, GTF2F2). Elevated DNA damage repair (DDR) activity in the G1 stage of the cell cycle aligns with the need for enhanced genomic surveillance before transitioning into the S phase, and was previously observed in proteomics experiments [72]. The Kinase ataxia-telangiectasia mutated (ATM) protein is an oncosuppressor Ser/Thr kinase integral part of the DDR pathways that is activated in response to DNA double strand breaks (DSBs). Its altered activity was observed in the nuclear fractions by both area and PSM measurements (with only PSM measurements passing the statistical filtering criteria). It acts by phosphorylating cell cycle checkpoint control and apoptotic response proteins resulting in cell cycle arrest at the G1/S, S and G2/M checkpoints [73]. Mutations in the ATM gene were shown to sensitize cancer to Pt-derived drugs, on one hand, but also to increased risk of secondary tumors after radiotherapy, on the other hand [74]. As cancer therapeutic drugs can induce DNA lesions, the DDR machinery is often activated in response to the treatment. Such a reaction has been associated with the emergence of both hypersensitivity and resistance to therapeutic agents, pointing to new cancer targeting opportunities that focus on the DNA repair pathways [75].

Altogether, the chromatin remodeling and transcription regulation processes were affected as evidenced by ATRX [76,77] and a number of zinc finger and histone proteins that upon phosphorylation, acetylation or ion binding have altered DNA binding and chromatin association ability (**Figure 8B**). It was encouraging to observe the upregulation of the PDCD4 protein (Programmed Cell Death 4 or Neoplastic Transformation Inhibitor) which is a tumor suppressor that inhibits translation initiation and suppresses tumor progression (**Figure 8B**). The expression level of this protein is often downregulated in breast cancers [78–81]. HER2 activation downregulates PDCD4 through the MAPK and PI3K/AKT pathways [79], and overexpression of PDCD4 was reported to promote apoptosis in breast cancer cell lines in the presence of the HER2 inhibitor trastuzumab [79,80]. The exact mechanism of PDCD4 inhibition of proliferation remains under research, but it has been suggested that PDCD4 inhibits translation by binding the translation initiation factor EIF4A [79,81]. Here, the change in abundance was detected in both cellular fractions.

In the cytoplasm, distinct processes that were upregulated as a whole pertained mostly to mitochondrial bioenergetics that encompassed proteins involved in mitochondrial organization, gene expression, electron transport and cellular respiration such as ATP synthases, cytochrome oxidases, oxidoreductases, and mitochondrial ribosomal proteins (MRPs) (**Figures 4, 8**, and **Supplemental File 3**). The upregulation of MTIF2 (Mitochondrial Translational Initiation Factor 2) and PDCD4 were additional indicators of inhibited cell proliferation. A consistent upregulation backed by both area and PSM measurements was also recorded for GADD45GIP1 (Growth Arrest and DNA Damage-Inducible Proteins-Interacting Protein 1) - an inhibitor of G1 to S cell cycle progression. GADD45GIP1 is a nuclear protein that can be also found in the mitochondria [21], and while its abundance was larger in the nuclear fraction, upregulation in the nuclear fraction was not observed. The accumulation of EGFR in the cytoplasm was noted consistently and was in agreement with prior findings that emerged from the treatment of various EGFR mutant cancers with Tyr kinase inhibitors where the cytoplasmic accumulation of EGFR was used as a predictor of clinical efficacy [82].

### Lapatinib/Ipatasertib treatment vs EGF stimulation of cells

Based on GO annotations, as many as 25 proteins could be mapped to the EGFR/ERBB2 and PI3K/PKB signal transduction and regulation pathways, several being shared by both (**Figure 7B**). PI3K/PKB is activated downstream of EGFR/ERBB2, hence, the two pathways collectively affect numerous downstream signaling processes by imposing redundant, additive or synergistic effects [83]. The effect of Ipatasertib was not always fully conclusive as the change was detected either by area or PSM measurements (but not both), or the statistical filters missed the pre-set thresholds, calling therefore for increased scrutiny prior to interpretation. Many of the PI3K/PKB components were already altered by the Lapatinib treatment alone (**Supplemental file 4**). Nonetheless, the impact of adding Ipatasertib to Lapatinib was observed on several proteins involved in the regulation of signal transduction (GRB2, PP2CA, LRP2, RRAS, DUSP3) (**Figure 7B**). Moreover, unique changes induced by the addition of Ipatasertib were reflected by alterations in the expression and/or activity of a number of proteins implicated in regulating gene expression (transcription factors TAF2, NCOR2, ZNF217, DRAP1), translation (CAPRIN1, GARS1), nuclear import/export (KPNA2, IPO7), protein trafficking (MYO1D, STXBP3), protein folding (PFDN2), DNA repair (RIF1, CDK5RAP3), cell cycle/growth (RHOA, RRAS2, CDK5RAP3), splicing (DDX5), transport (ABCC1 or MRP1 - Multidrug resistance-associated protein 1), and several metabolic, mitochondrial and biosynthetic processes [21] (**Figure 9A**). Processes related to the “downregulation of the negative regulation of apoptotic signaling pathway” (YBX3, HMGB2, CTNNA1, HTT, GCLM, HSPA1B, DNAJA1, CD44) and “protein folding” were impacted to a larger extent in the Lapatinib/Ipatasertib-treated cells (**Figure 4D**). Some of the affected proteins have raised interest in prior studies. Among these were the transcriptional repressors NCOR2 (a nuclear receptor co-repressor) and DRAP1 (involved in the repression of class II genes transcription), and ABCC1/MRP1 - an ABC transporter that mediates the efflux of drugs from cells. The downregulation of GDI1 (nuclear fraction, PSM measurements only) and TUBB3 (cytoplasmic fraction, area measurements only) also appeared to be affected to a larger extent by the combination of drug treatments. GDI1 (GDP Dissociation Inhibitor 1) has been shown to act as both tumor suppressor and oncogene, up- or downregulation being dependent on the cancer type [84–87]. Its overexpression in colorectal and hepatocellular cancer cells was found to lead to telomere dysfunction, and cell proliferation and migration, respectively, through the PI3K/AKT pathway [86,87]. TUBB3 response to PI3K/AKT pathway inhibitors was also reported to be either upregulated or downregulated in various cancer types [88–91].

### Other affected categories

The drug treatments affected several additional proteins that were previously associated with cancer and metastatic processes, however, these proteins are still in early stages of investigation with limited information regarding their function and significance. For example, emerging evidence has uncovered tumor promoting (via involvement in immune escape, angiogenesis, EMT) or suppressive (via participation in growth and proliferation) roles for several solute carriers (SLCs) [92]. The altered metabolism of cancer cells is characterized by an increased uptake of glucose and lactate released by the cells of the TME. Recent reviews have highlighted specific roles for the SLC families involved in the uptake of glucose (SLC2A), lactate (SLC16A), glutamine (SLC1A5), leucine (SLC7A5/SLC3A2), and cystines (SLC7A11) [92]. In this work, most SLCs did not change abundance but some displayed some level of downregulation, among which the glucose SLC2A1 uptake transporter (**Figure 8A**). While some of the SLCs were detectable in both nuclear and cytoplasmic fractions, it was intriguing that the abundance of some carriers appeared to be higher in the nuclear fractions (especially of SLC2A1) where the observed changes were measured.

CRYBG1 (suppression of tumorigenicity 4) is a protein involved in cytoskeletal remodeling with potential roles in suppressing melanoma [21] that was downregulated in both nuclear and cytoplasmic cell fractions (**Figure 8A**). Its loss of function, however, was related to enhanced invasive and migratory capabilities in cancer cells [21,93]. The expression of the Ser protease ST14 (Tumor associated differentially expressed gene 15 protein or Suppressor of tumorigenicity 14 protein) has been also associated with cancerous states [21], and its overexpression in early ovarian cancers was correlated with longer survival rates suggesting its potential as a prognostic marker [94]. It was also found to reduce cell migration [94]. Here, the protein exhibited a decrease in expression in the cytoplasmic fractions following the drug treatment, confirmed by fold-changes according to both area and PSM measurements, but only with the area measurements passing the statistical filtering criteria.

Finally, several key elements including oncogenes, tumor suppressors, angiogenic/EMT metastatic factors, and epithelial/mesenchymal markers directly linked to cancer progression and detection (**Figure 9B**). The downregulation of oncogenes, EMT regulators and of angiogenic factors, coupled with the upregulation of tumor suppressor proteins, were all indicators of positive outcomes for suppressing aberrant proliferation. The upregulation of DAG1 which is involved in the establishment of cell-ECM interactions, was shown to impair EMT processes [95]. Additionally, the upregulation of epithelial- and downregulation of mesenchymal markers are well-established indicators of reduced migration and invasion capabilities [96], in this case attributed to the drug treatments.

### Potential drug targets

Many of the detected proteins represent approved drug targets (27 proteins) in the DrugBank database, and yet many more (42 proteins) are investigational targets (**Figure 10**). The up- or downregulated proteins in either the nuclear or cytoplasmic fractions constitute an emerging pool of potentially novel targets, with several being already proposed for therapeutic intervention, for example: SRPK1 (Serine/Arginine-Rich Splicing Factor Kinase 1) (downregulated in the nuclear and cytoplasmic fractions) - a protein involved in the regulation of splicing and whose inhibition would simultaneously affect diverse and distinct processes such as migration, metastasis, apoptosis, and sensitivity to chemotherapy in breast cancer [97]; PTPRD (upregulated in the cytoplasmic fractions) - a tyrosine phosphatase tumor suppressor that is implicated in cell growth inhibition [98,99]; and AGRN (downregulated in the nuclear and cytoplasmic fractions) - a glycoprotein that promotes cell growth and invasion through the PI3K/AKT pathway, and immune cell infiltration [100]. VTCN1 - a cell membrane protein that inhibits T-cell responses and for which studies have shown that increased expression promotes cancer cell proliferation [21,101], was, interestingly, upregulated in the cytoplasmic fractions in response to the drug treatments, raising thus interest for exploration in the context of immunotherapies. The downregulation of proteins such as ANLN and endophilin A2 uncovered additional candidates for the development of synergistic therapeutic strategies with increased effectiveness. Endophilin A2 has been previously associated with metastatic processes due to its roles in endocytosis, cytoskeletal dynamics, cell migration and cancer cell invasion [21,67]. Mitochondrial ribosomal proteins (MRPs) have important roles in mitochondrial translation and protein synthesis necessary for oxidative phosphorylation, cellular respiration, and energy production. Their abnormal expression has been reported is several cancers, including breast cancer, and was linked to apoptosis and cell death initiation, tumorigenesis, and other processes that derive from the complexity and variety of the differentially expressed MRPs. Taken altogether, mitochondrial dysfunction and apoptosis constitute hallmarks of cancer that are of importance to emerging targeted interventions [102–104]. Several MRPs that were upregulated upon drug treatment are known to promote apoptosis and present potential interest. These included the death-associated protein 3 (DAP3/MRPS29) - an AKT substrate that binds to apoptotic factors and TRAIL receptors [102], MRPS30 - which induces apoptosis independent of the death receptor-induced extrinsic pathway [103], and MRPL11 - which indirectly activates p53 and leads to cell cycle arrest [105].

## Conclusions

Proteomic profiling of HER2+ breast cancer cells treated with Lapatinib and Ipatasertib was performed for the first time to provide a comprehensive view of their combined impact on cellular biology and shed light on the mechanisms of drug action and possible off-target effects. Complementary to the inhibition of EGFR/ERBB2 and PI3K/AKT pathways, the two drugs affected essentially all cancer hallmark processes, downregulating cell cycle progression, adhesion, immune response and inflammation, while upregulating chromosome organization, transcription, DNA damage repair, and mitochondrial bioenergetics. Ipatasertib further altered the expression or activity of proteins involved in transcription, cell cycle, protein transport and trafficking, and downstream signaling pathways. Changes in the expression of key cancer drivers such as oncogenes, tumor suppressors, EMT and angiogenesis regulators underscored the inhibitory effectiveness of drugs on cancer proliferation. Diagnostics and multidrug resistance markers emerged, as well, revealing intrinsic resources for assessing the cellular response to drug inhibition and predicting metastatic potential. Overall, this study provides valuable insights into the molecular effects of Lapatinib and Ipatasertib kinase inhibitors, expanding the opportunities for the development of novel and combinatorial targeted therapies.

## Data availability

The mass spectrometry raw files were deposited to the ProteomeXchange Consortium via the PRIDE [106] partner repository with the dataset identifier: PXD051094.

## Author contributions

AK performed the experiments; NRM acquired the preliminary data; AK and IML analyzed the data and wrote the manuscript; IML coordinated the work. All authors reviewed and approved the final version of the manuscript.

## Acknowledgment

This work was supported by an award from the National Institute of General Medical Sciences (Grant No. 1R01GM121920) to IML.

