## Supplemental file 1 for "Proteomic Assessment of SKBR3/HER2+ Breast Cancer Cellular Response to Lapatinib and Investigational Ipatasertib Kinase Inhibitors"

**PSM detection reproducibility.** Scatter plots representing the reproducibility of PSMs in three biological replicates of the nuclear and cytoplasmic fractions for each drug treatment experiment.

#### PSMs reproducibility (Nuclear fractions)

| EGF | Lapatinib | Lapatinib<br>Ipatasertib |
| --- | --- | --- |

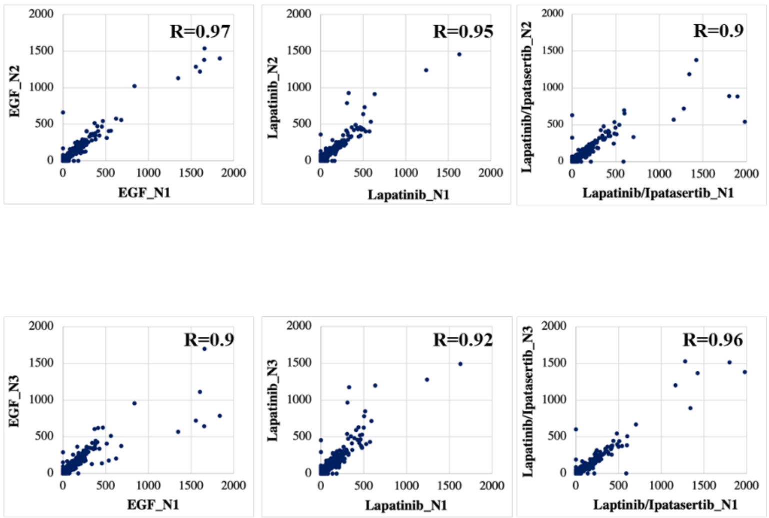


#### PSMs reproducibility (Cytoplasmic fractions)

| EGF | Lapatinib | Lapatinib<br>Ipatasertib |
| --- | --- | --- |

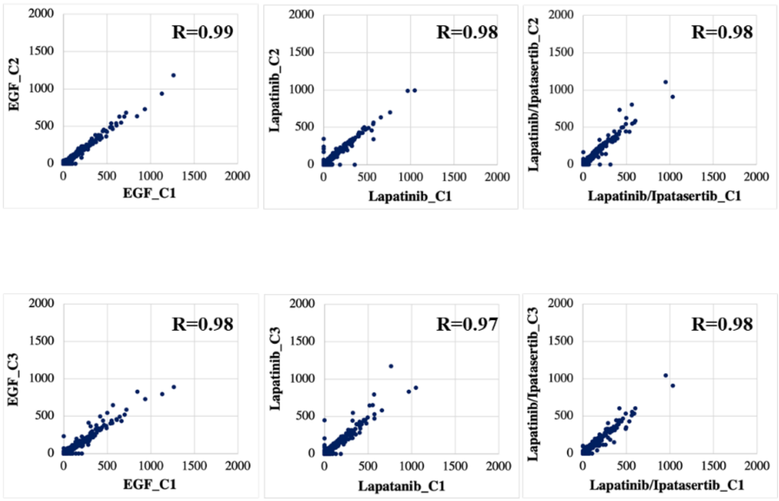

**Peptide detection reproducibility.** Scatter plots representing the reproducibility of peptide elution times and XCorr scores from each biological replicate of the cytoplasmic and nuclear fractions: replicate 1 (X-axis), replicate 2 (Y-axis), and replicate 3 (color bar).

### Peptide reproducibility (XCorr)

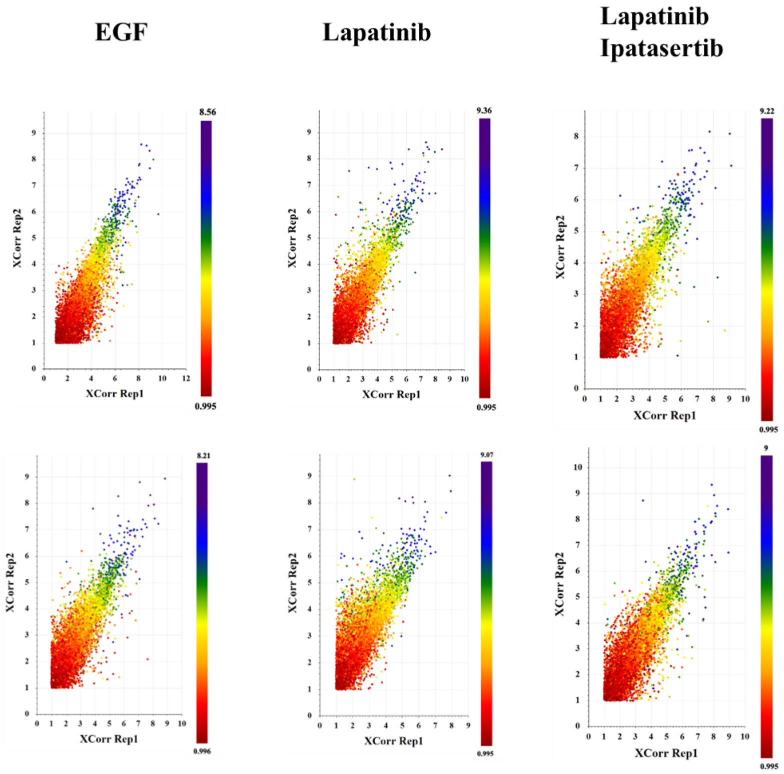


**Protein differential expression.** Volcano plots representing protein abundance measurements in the nuclear and cytoplasmic fractions of drug-treated Lapatinib/Ipatasertib vs Lapatinib-control cells. Differentially expressed proteins are indicated in red (up-regulated) and blue (down-regulated), displaying ≥2-fold change in abundance (p-value ≤0.05)

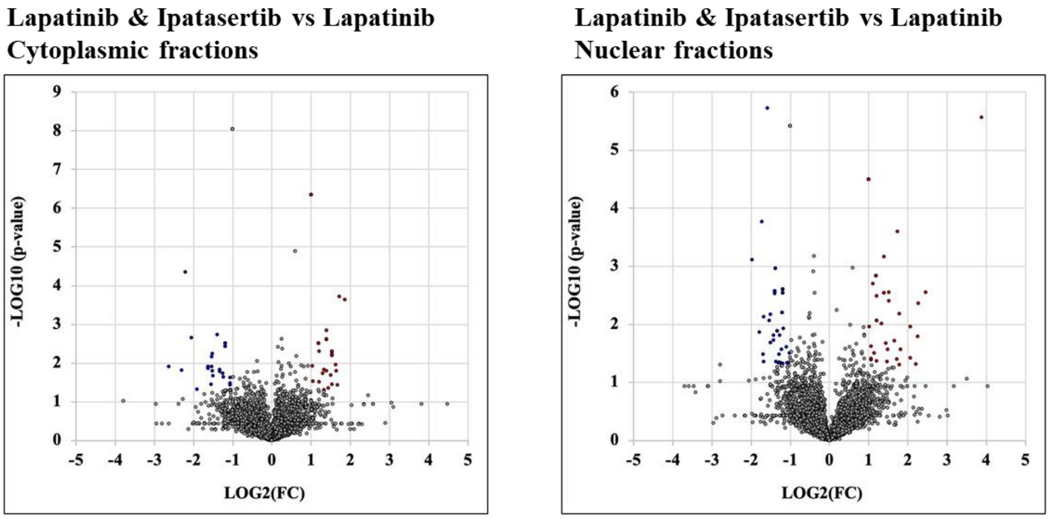
