## Supplemental file 6 for "Proteomic Assessment of SKBR3/HER2+ Breast Cancer Cellular Response to Lapatinib and Investigational Ipatasertib Kinase Inhibitors"

Parallel Reaction Monitoring (PRM)/MS validation of selected proteins that changed expression level in response to the drug treatments.

#### ***Upregulated in Lapatinib***

**Serine-protein kinase ATM (ATM)**
DIVDGMGITGVEGVFR, Charge +2, m/z = 832.9193 Da

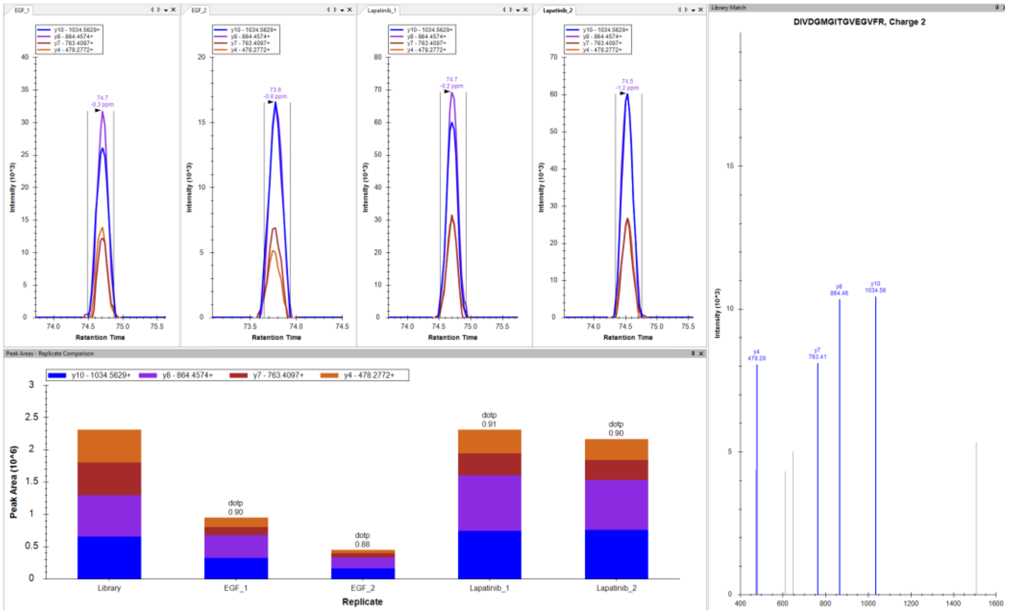


**Serine-protein kinase ATM (ATM)**
MTTDPMLAYLQPFR, Charge +2, m/z = 842.4153 Da

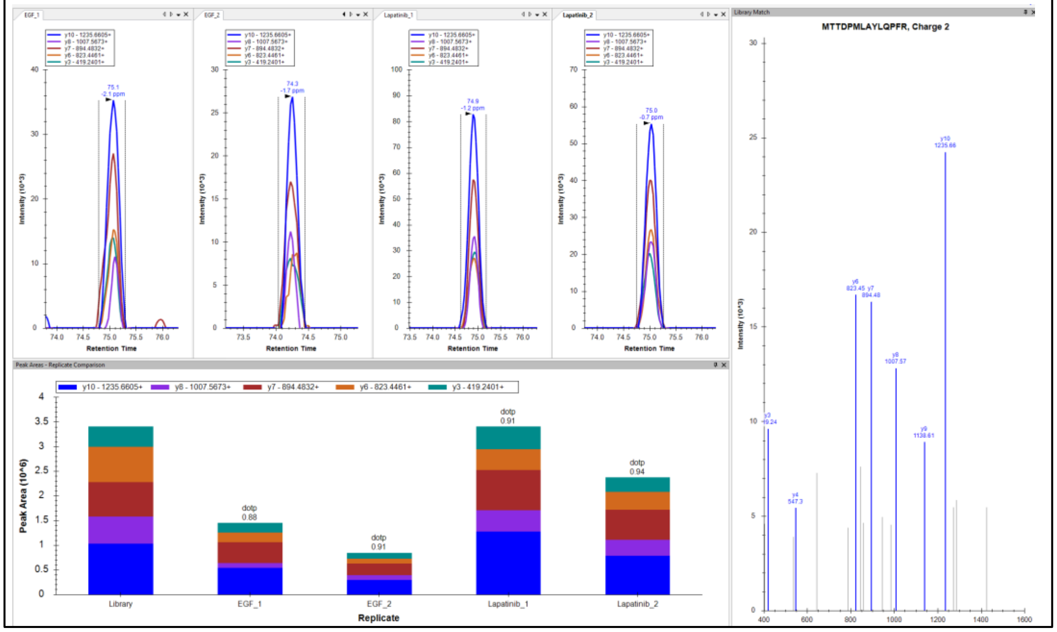

**Nuclear mitotic apparatus protein 1 (NUMA1)**
EFASHLQQLQDALNELTEEHSK, Charge +3, m/z = 856.4173 Da

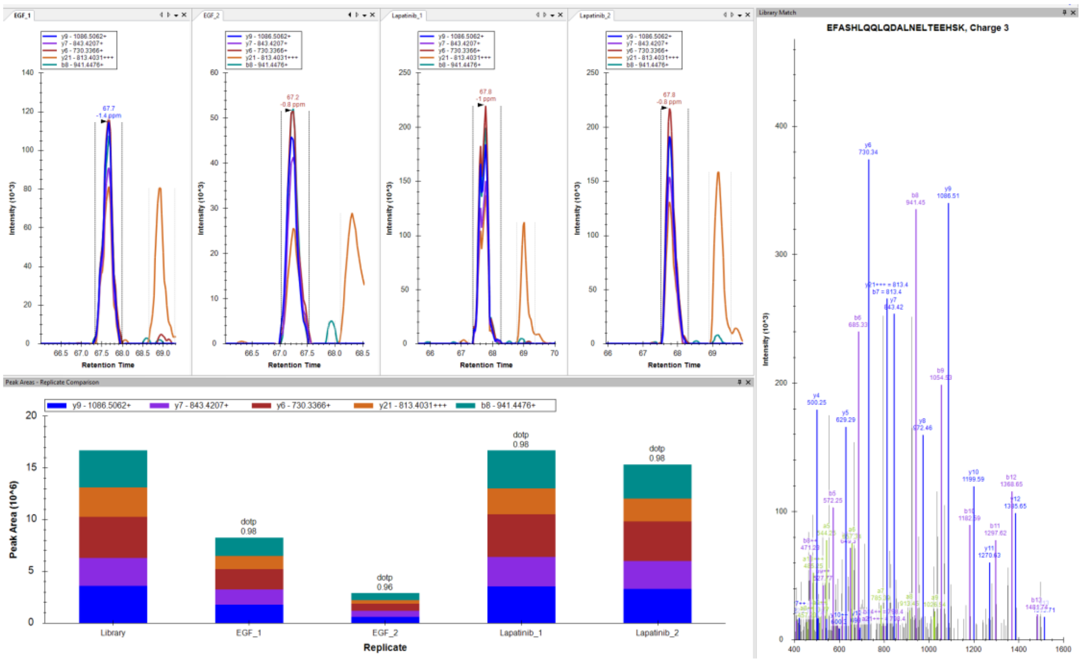


**Nuclear mitotic apparatus protein 1 (NUMA1)**
AQELGHSQSALASAQR, Charge +2, m/z = 827.4188 Da

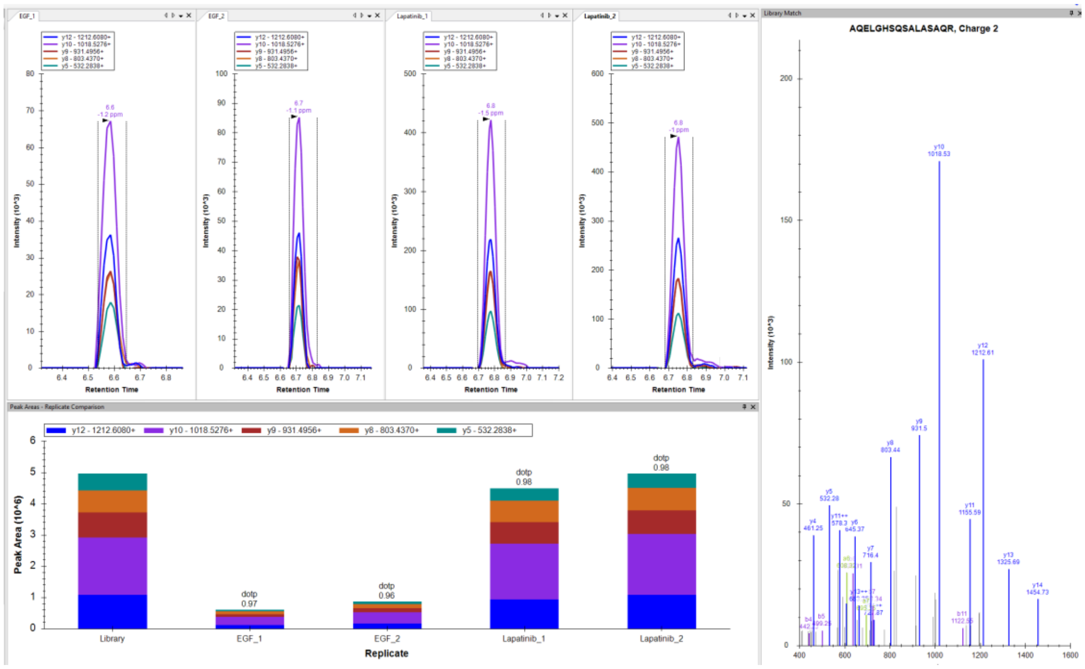

**Programmed cell death protein 4 (PDCD4)**
APQLVGQFIAR, Charge +2, m/z = 600.3484 Da

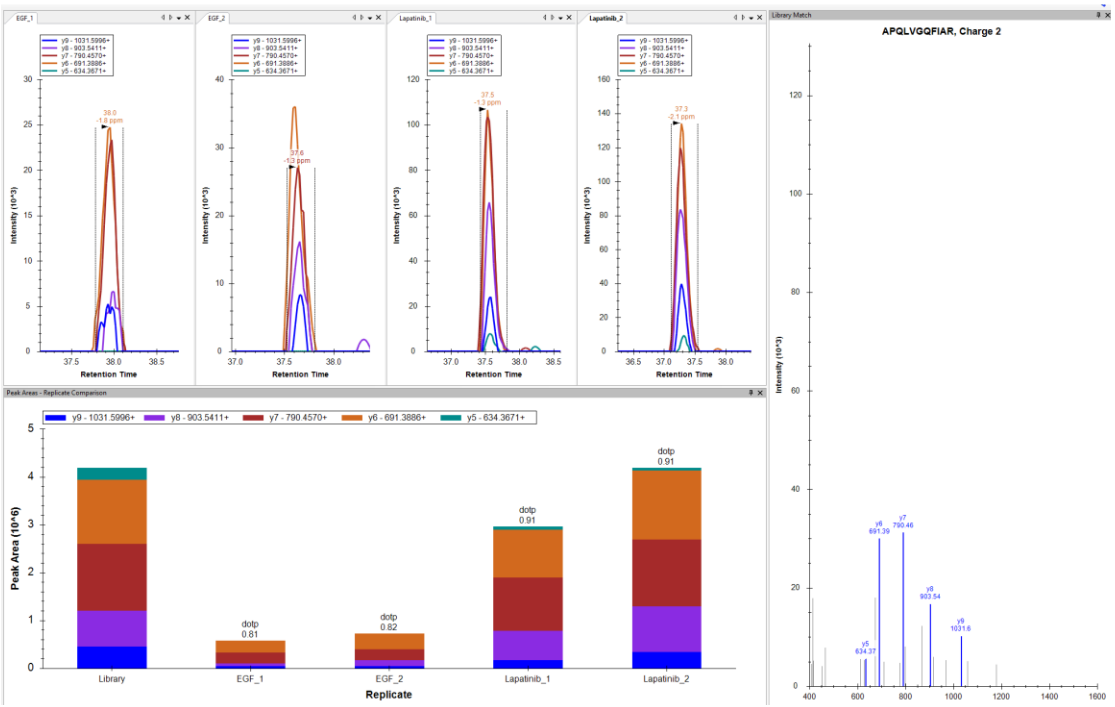


**Programmed cell death protein 4 (PDCD4)**
SGVPVLAVSLALEGK, Charge +2, m/z = 720.4270 Da

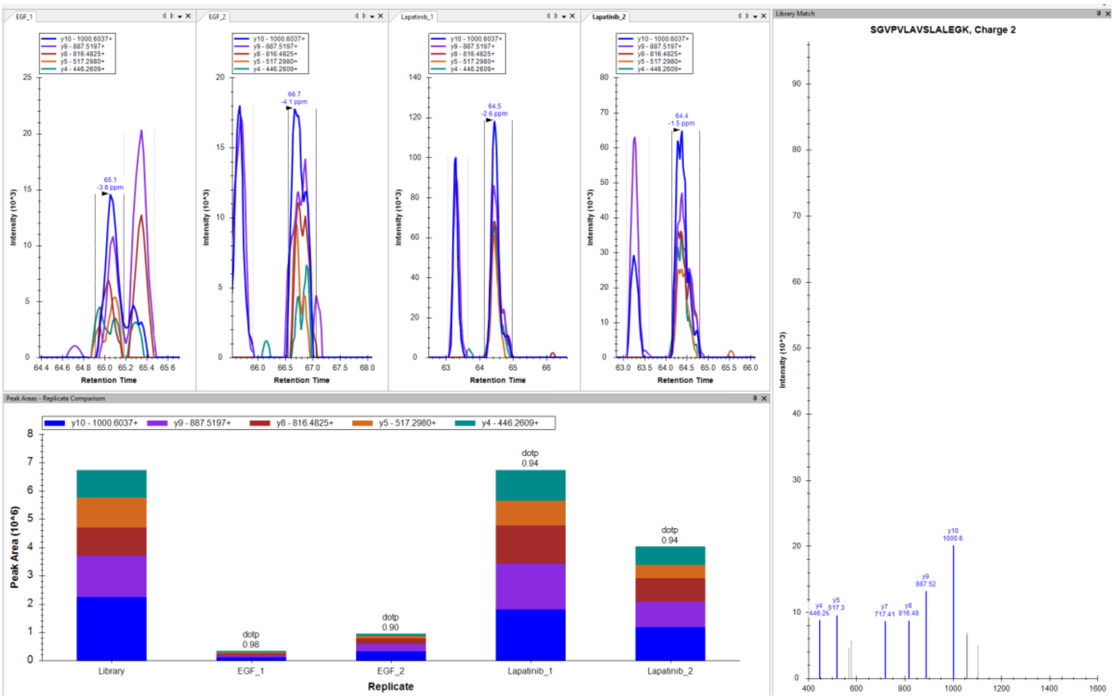

***Downregulated in Lapatinib***

**DNA topoisomerase 2-alpha (TOP2A)**
VTIDPENNLISIWNNGK, Charge +3, m/z = 643.0022 Da

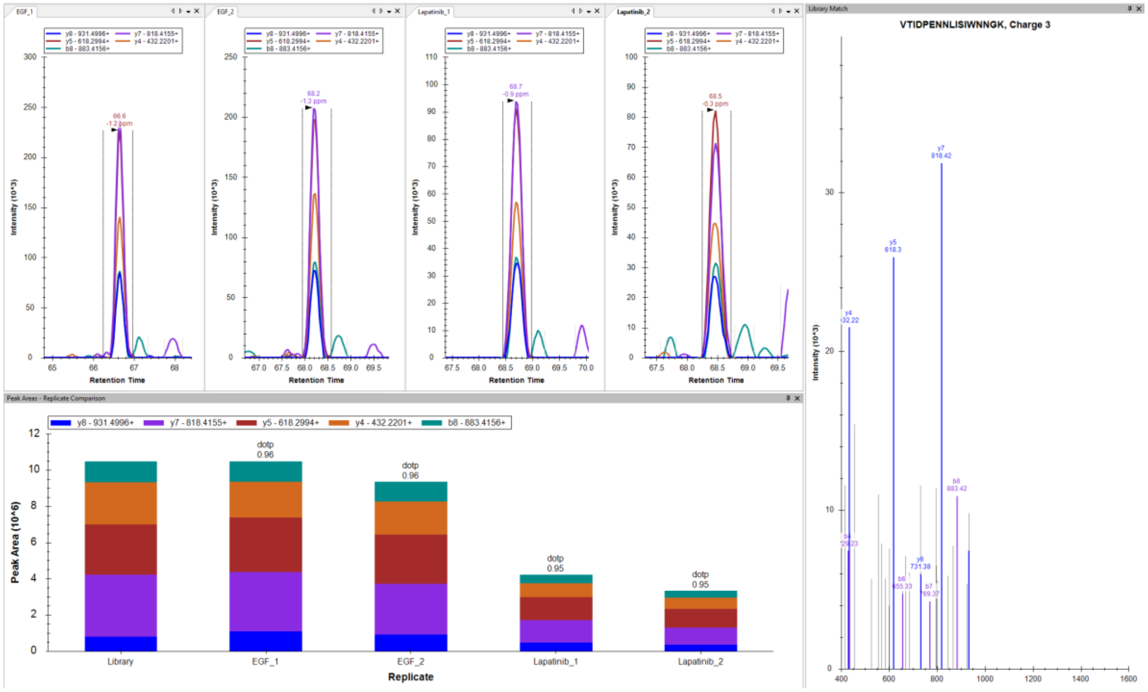


**DNA topoisomerase 2-alpha (TOP2A)**
EDLATFIEELEAVEAK, Charge +2, m/z = 903.9540 Da

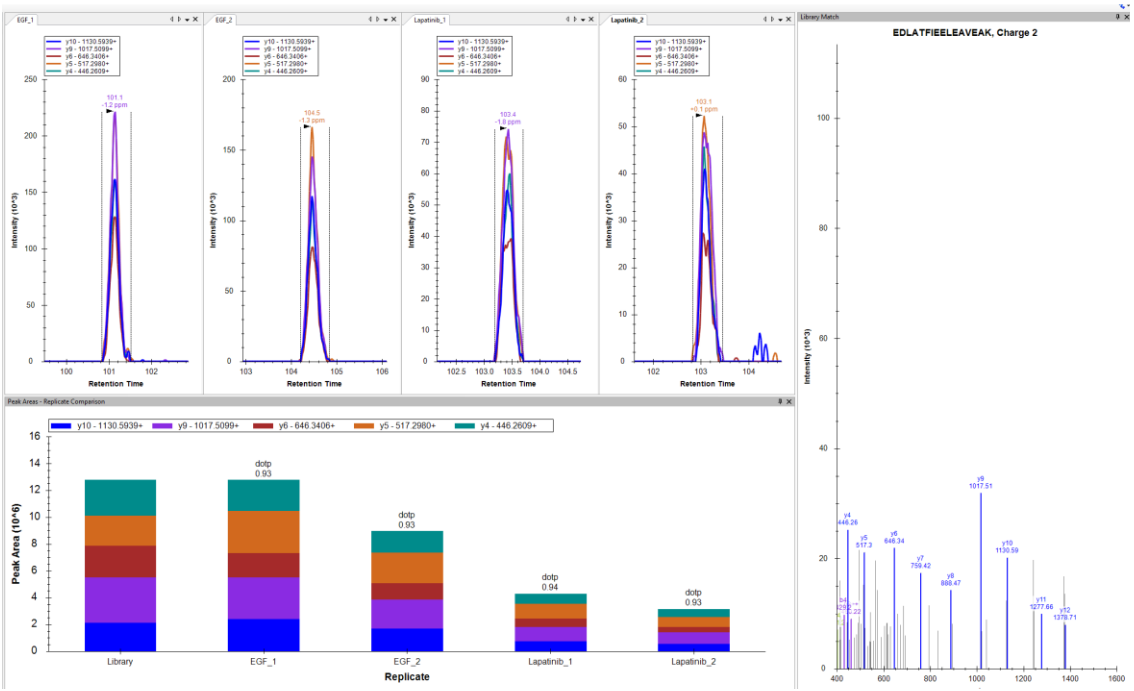

**Proliferation marker protein Ki-67 (MKI67)**
AVGASFPLYEPAK, Charge +2, m/z = 675.3586 Da

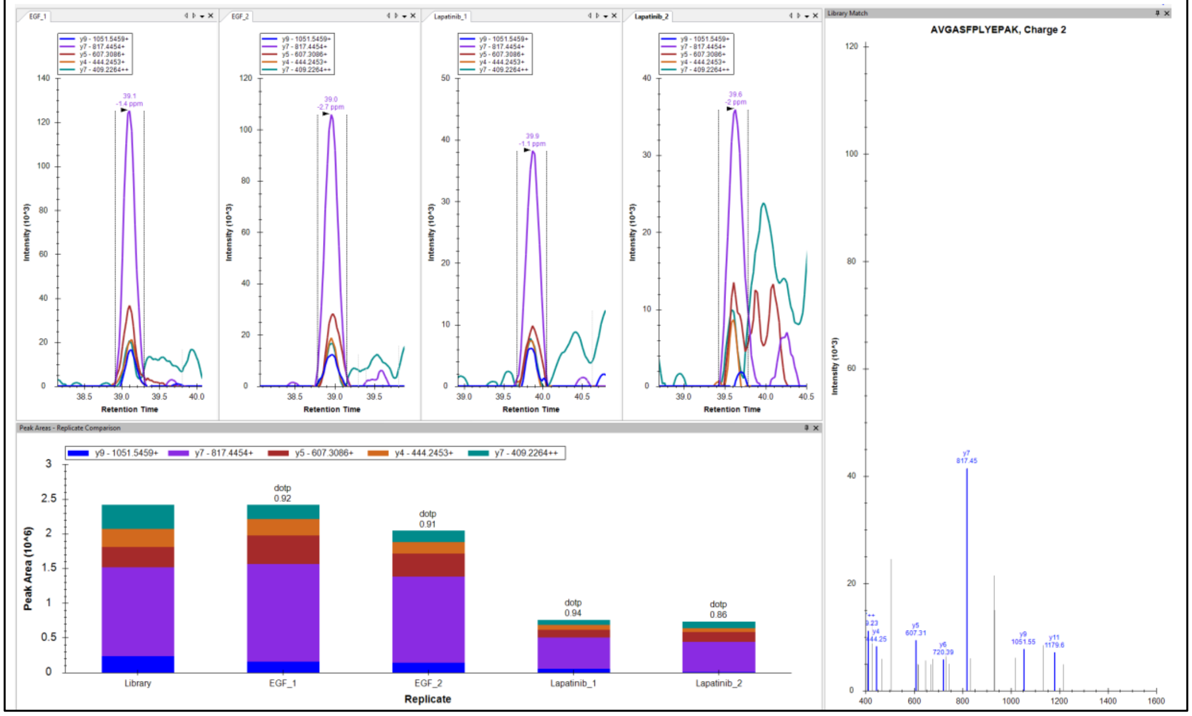

**Proliferation marker protein Ki-67 (MKI67)**
AQALEDLAGFK, Charge +2, m/z = 581.8088 Da

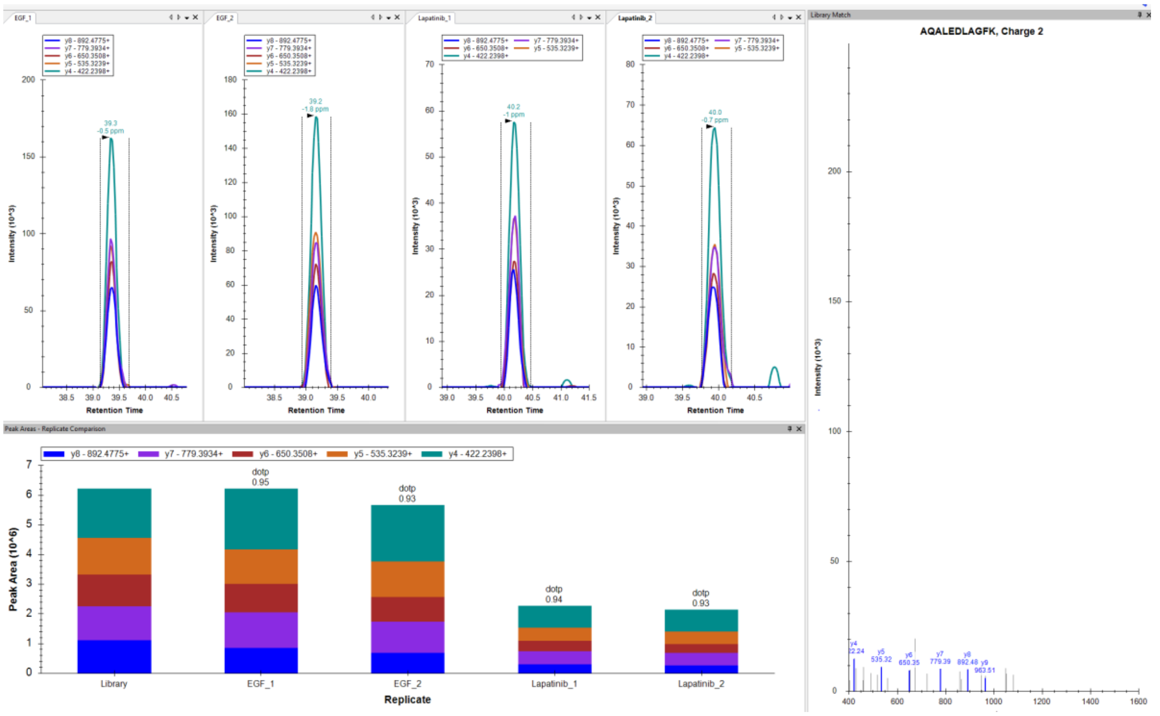

**14-3-3 protein sigma (SFN)**

VETELQGVCDTVLGLLDSHLIK, Charge +3, m/z = 794.7577 Da

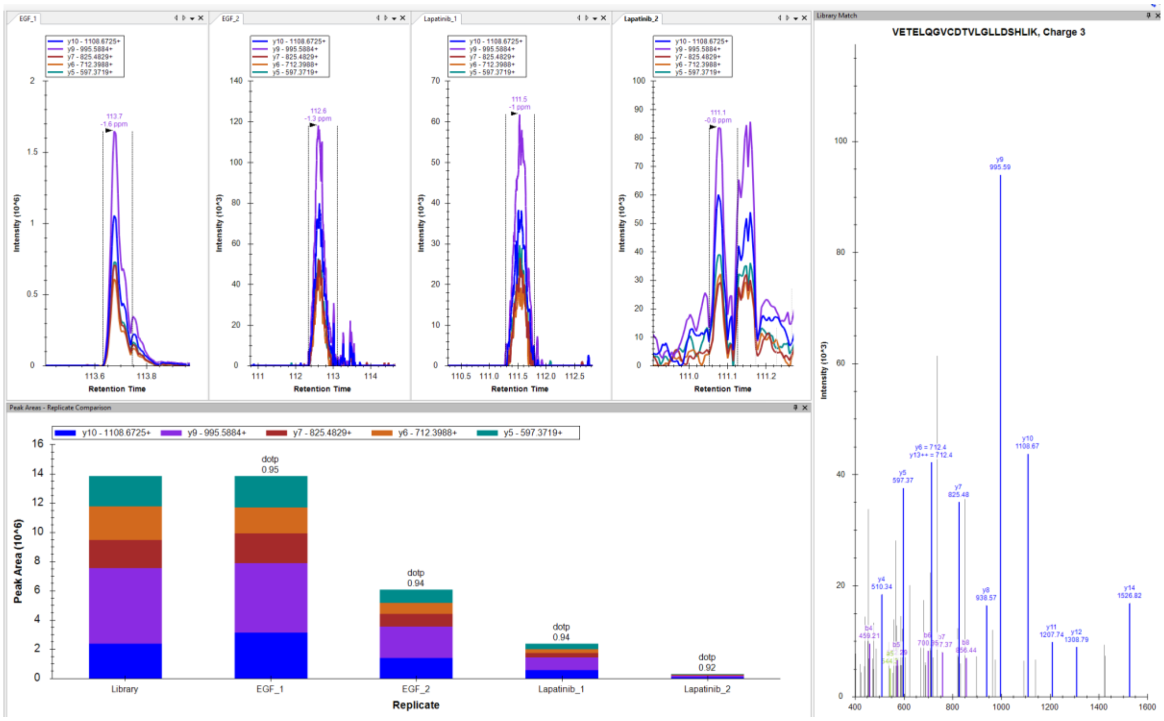


**CD44 antigen (CD44)**

YGFIEGHVVIPR, Charge +3, m/z = 462.9225 Da

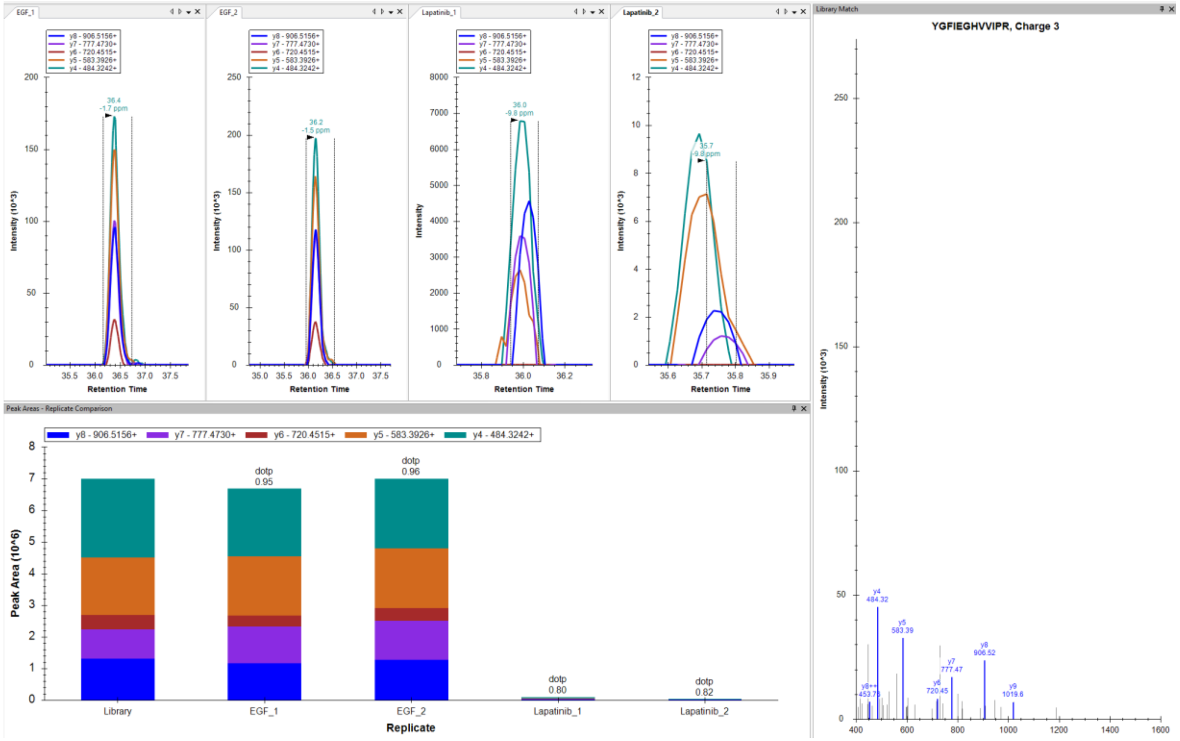

### ***Upregulated in Lapatinib/Ipatasertib***

**Serine-protein kinase ATM (ATM)**
MTTDPMLAYLQPFR, Charge +2, m/z = 842.4153 Da

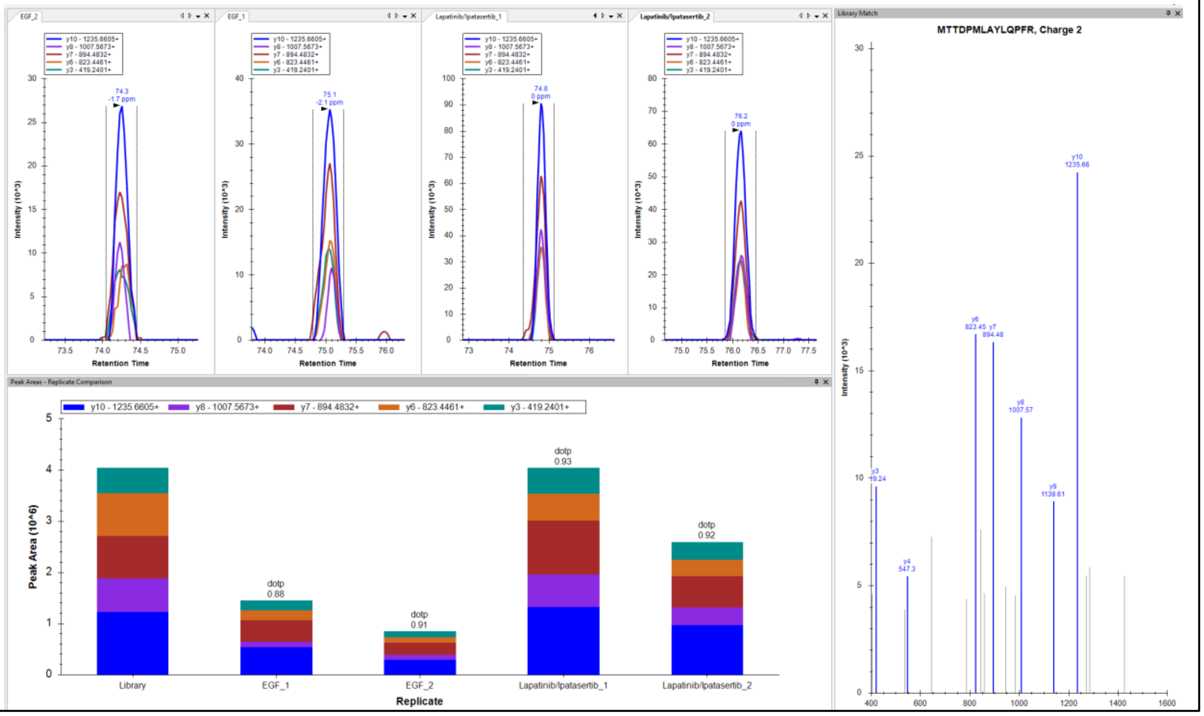


**Serine-protein kinase ATM (ATM)**
DIVDGMGITGVEGVFR, Charge +2, m/z = 832.9193 Da

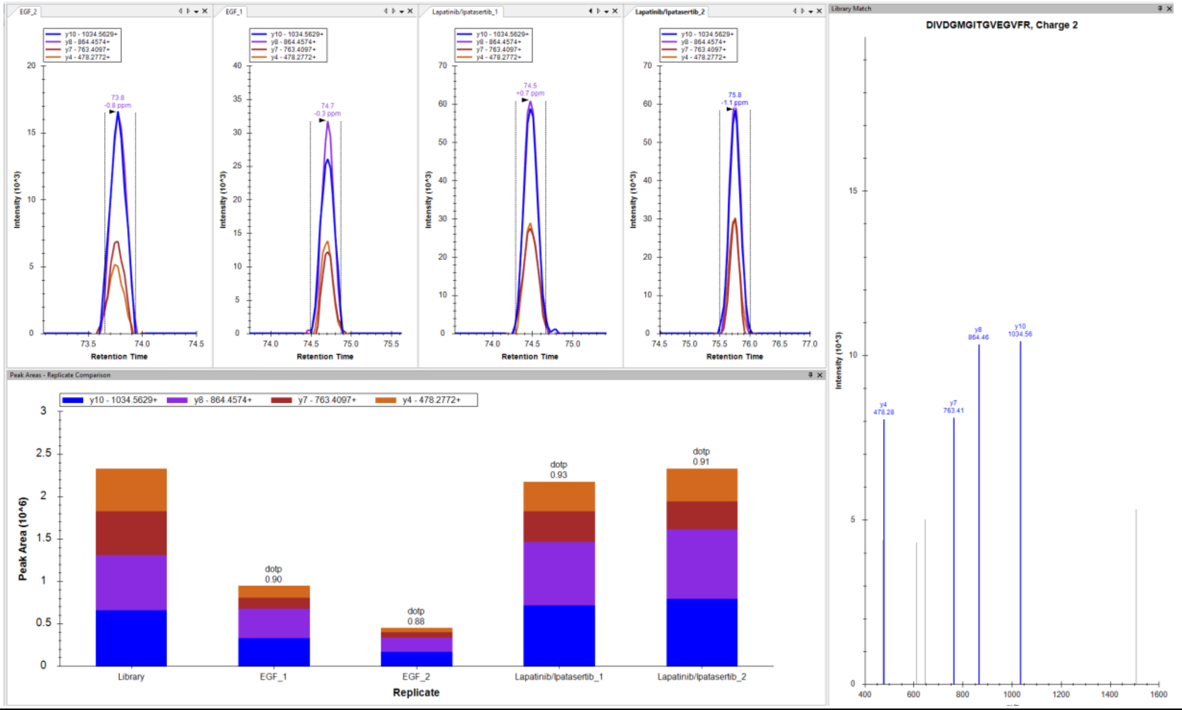

**Nuclear mitotic apparatus protein 1 (NUMA1)**
ASMQPIQIAEGTGITTR, Charge +2, m/z = 887.4618 Da

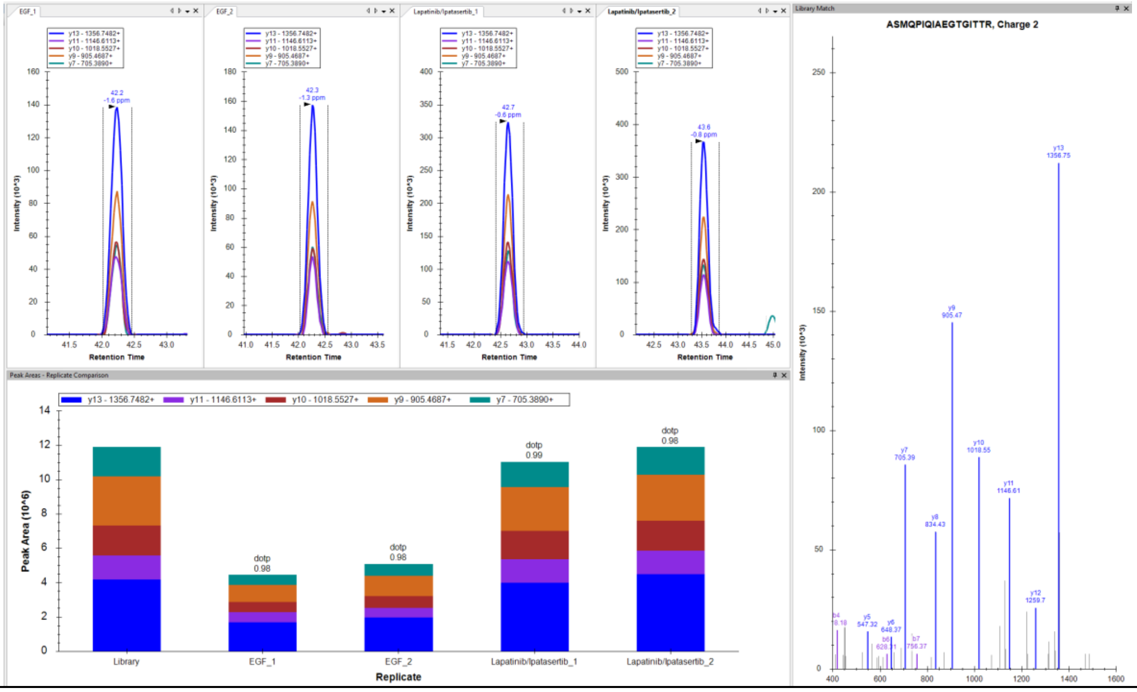


**Nuclear mitotic apparatus protein 1 (NUMA1)**
EFASHLQQLQDALNELTEEHSK, Charge +3, m/z = 856.4173 Da

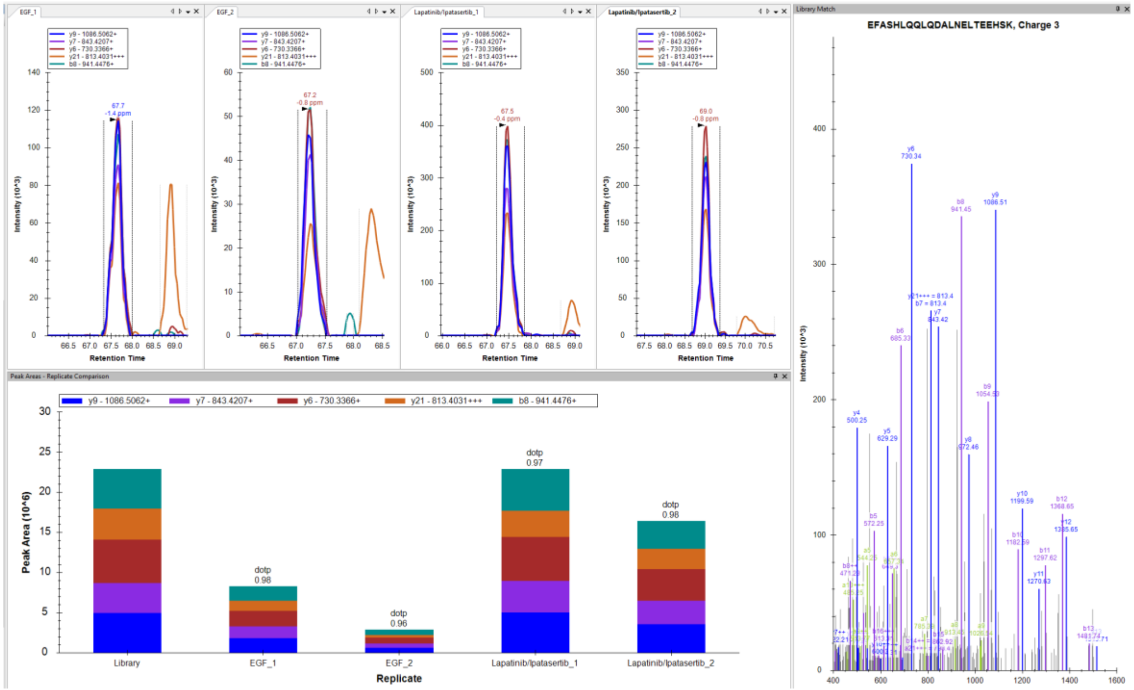

**Programmed cell death protein 4 (PDCD4)**
SGVPVLAVSLALEGK, Charge +2, m/z = 720.4270 Da

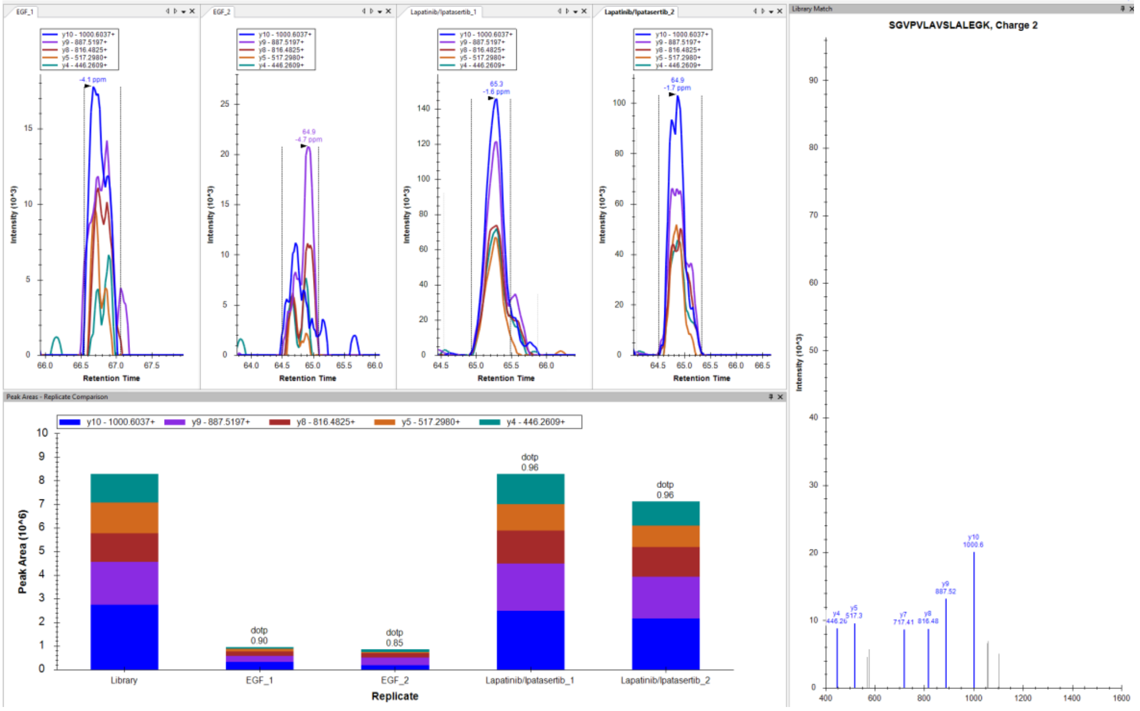


**Programmed cell death protein 4 (PDCD4)**
IYNEIPDINLDVPHSYSVLER, Charge +3, m/z = 829.4236 Da

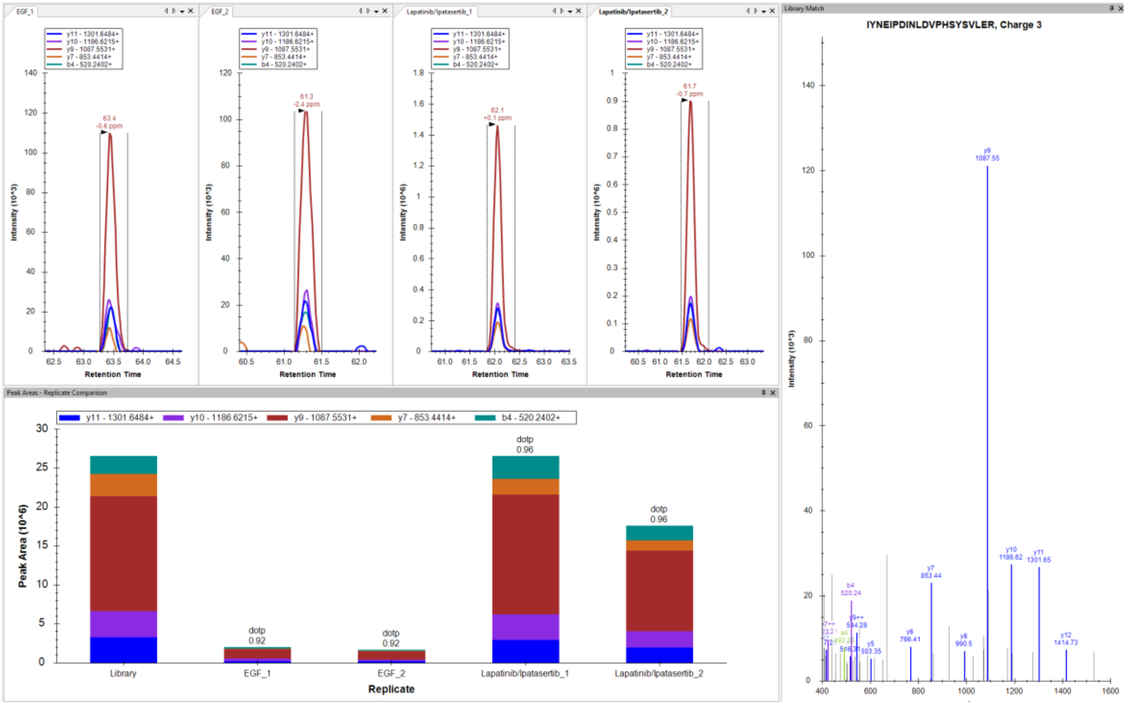

**V-set domain-containing T-cell activation inhibitor 1 (VTCN1)**
NVQLTDAGTYK, Charge +2, m/z = 605.3091 Da

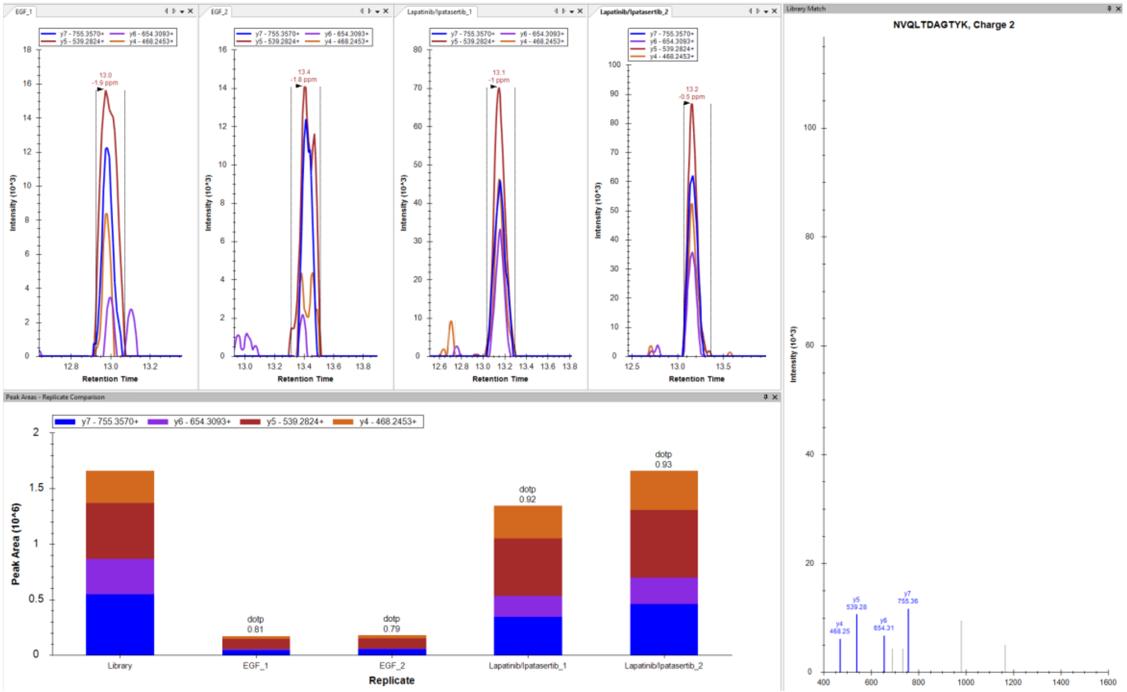


**V-set domain-containing T-cell activation inhibitor 1 (VTCN1)**
LSDIVIQWLK, Charge +2, m/z = 607.8608 Da

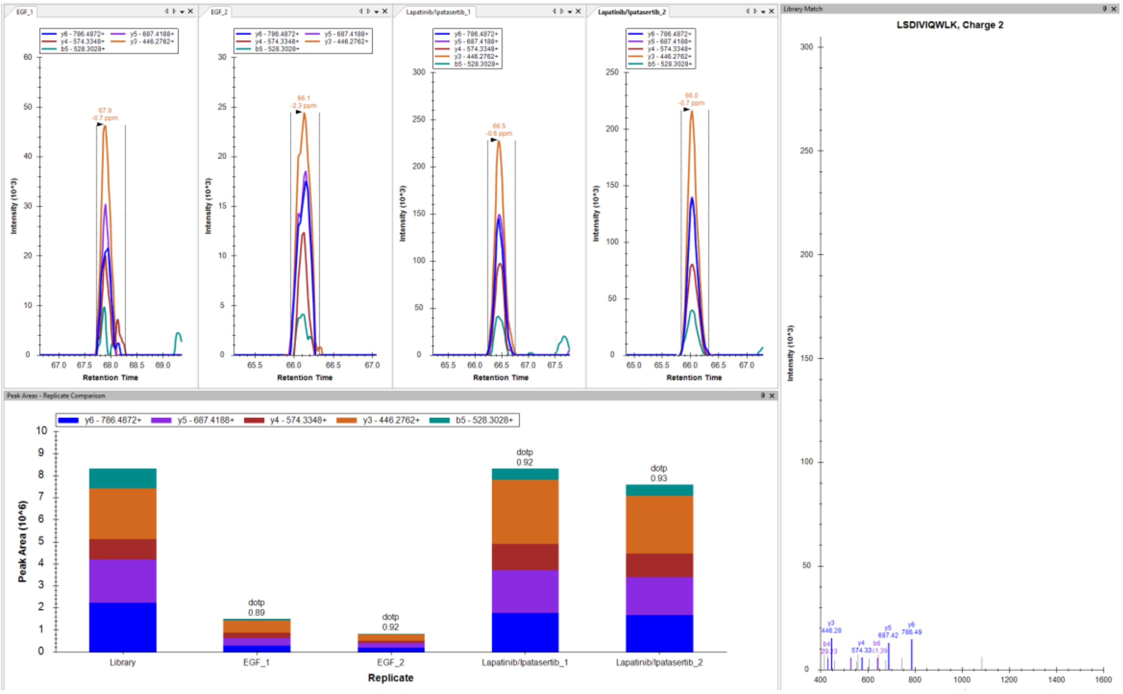

### ***Downregulated in Lapatinib/Ipatasertib***

**DNA topoisomerase 2-alpha (TOP2A)**
VTIDPENNLISIWNNGK, Charge +3, m/z = 643.0022 Da

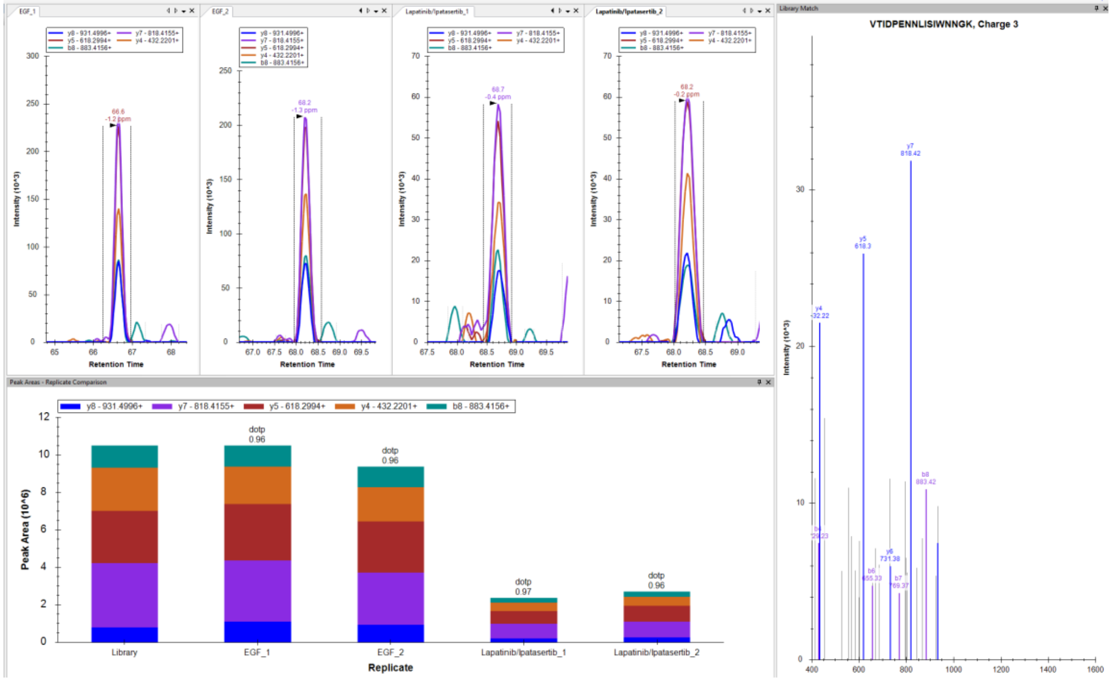


**DNA topoisomerase 2-alpha (TOP2A)**
EDLATFIEELEAVEAK, Charge +2, m/z = 903.9540 Da

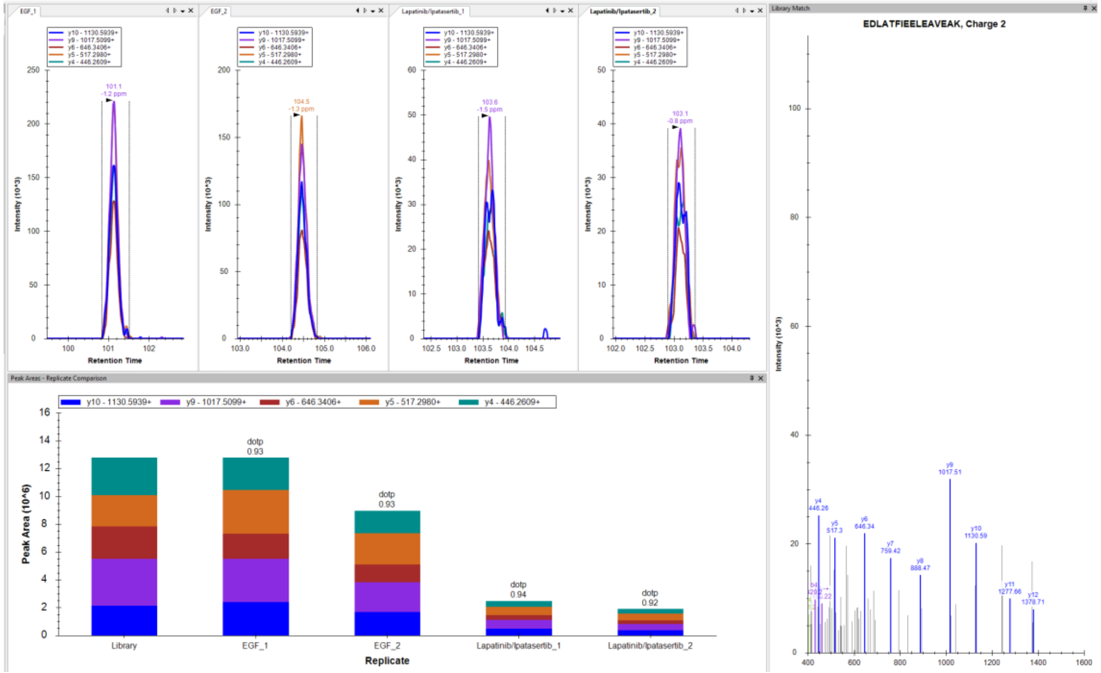

**Proliferation marker protein Ki-67 (MKI67)**
AQALEDLAGFK, Charge +2, m/z = 581.8088 Da

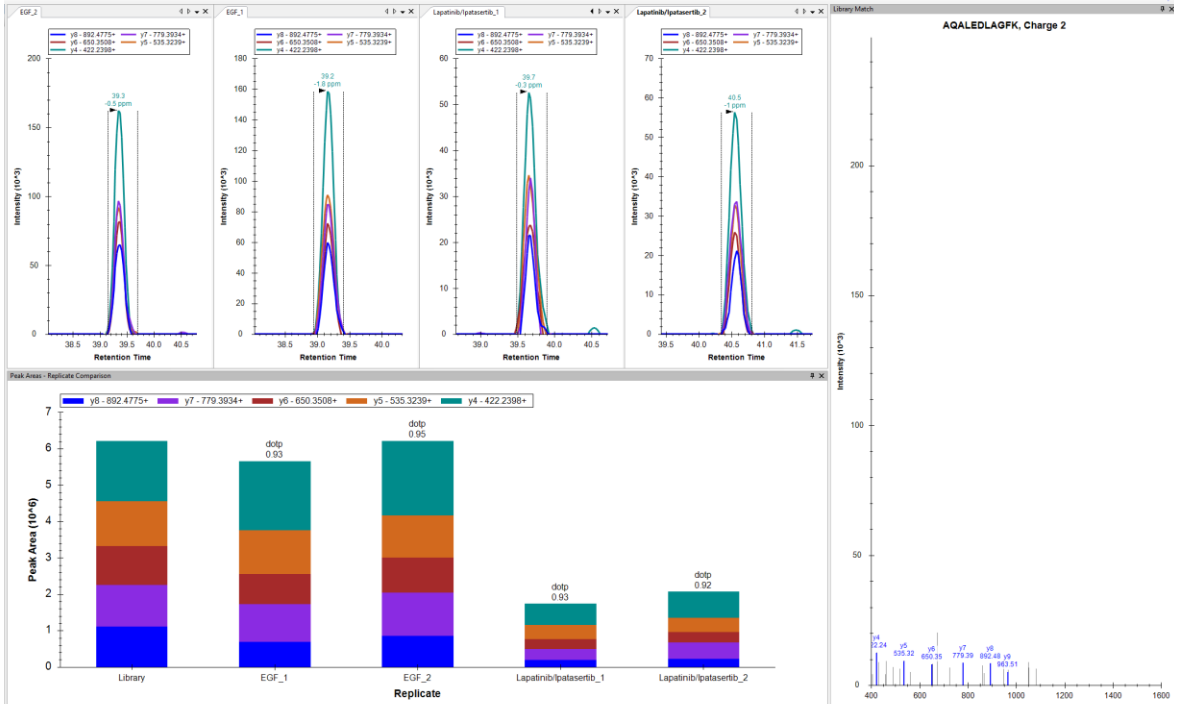


**Proliferation marker protein Ki-67 (MKI67)**
AVGASFPLYEPAK, Charge +2, m/z = 675.3586 Da

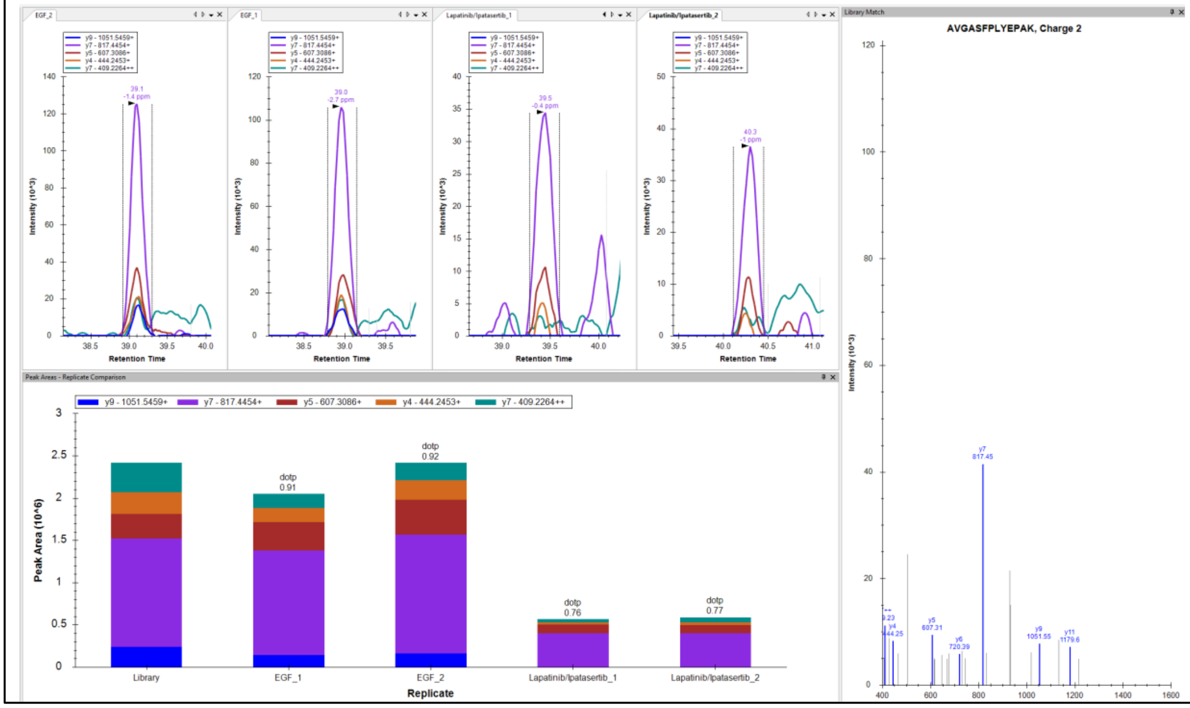

**14-3-3 protein sigma (SFN)**

VETELQGVCDTVLGLLDSHLIK, Charge +3, m/z = 794.7577 Da

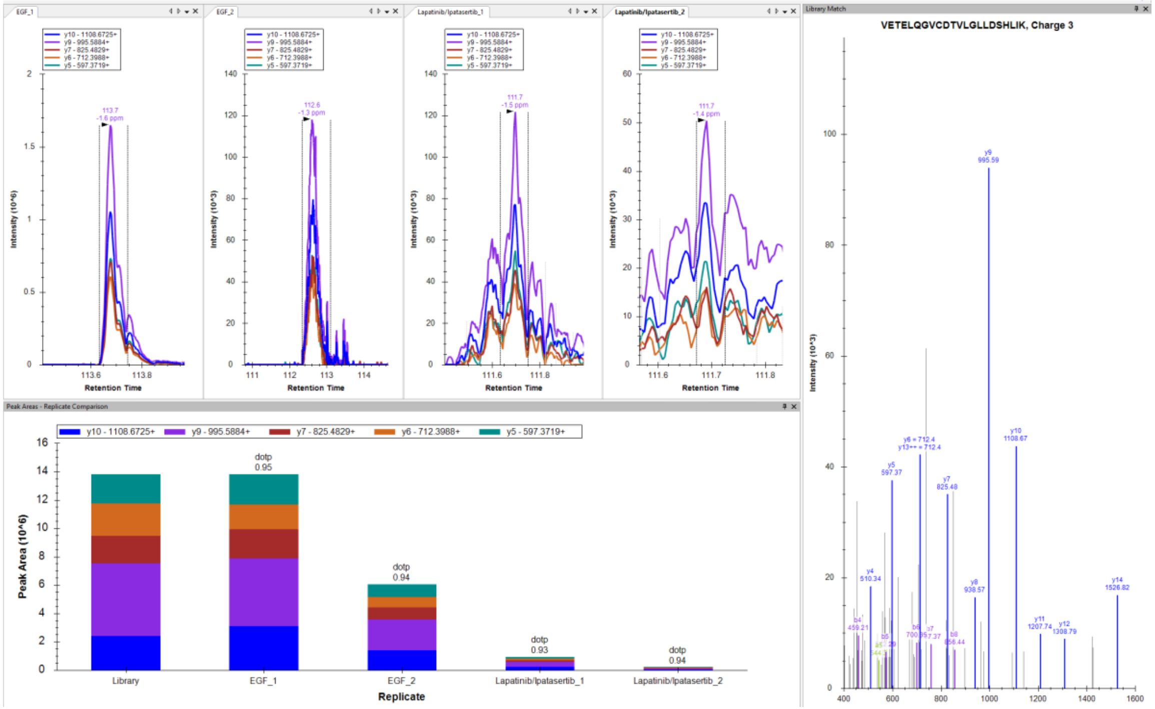


**CD44 antigen (CD44)**

YGFIEGHVVIPR, Charge +3, m/z = 462.9225 Da

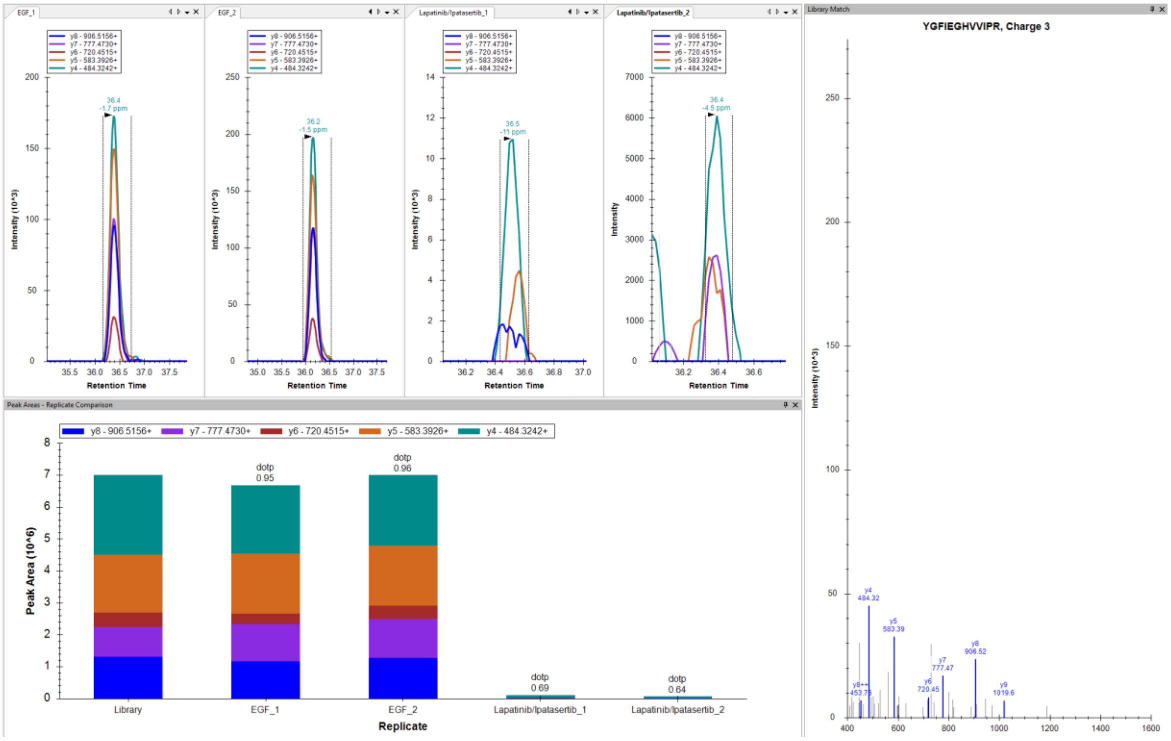
