## Supplemental file 7 for "Proteomic Assessment of SKBR3/HER2+ Breast Cancer Cellular Response to Lapatinib and Investigational Ipatasertib Kinase Inhibitors"

Western blot validation of selected proteins that changed expression level in response to the drug treatments.

**Protein:** VTCN1/ B7-H4/ Q7Z7D3, MW = 75 kDa, upregulated in L vs EGF and L/I vs EGF (cytoplasmic fraction)

**Conditions:**

- Primary Ab –VTCN1 (CST-14572) 1:1000
- Secondary Ab – Anti Rabbit HRP conjugate (CST- 7074S) 1:2000

*Note: L-lapatinib, I-ipatasertib*

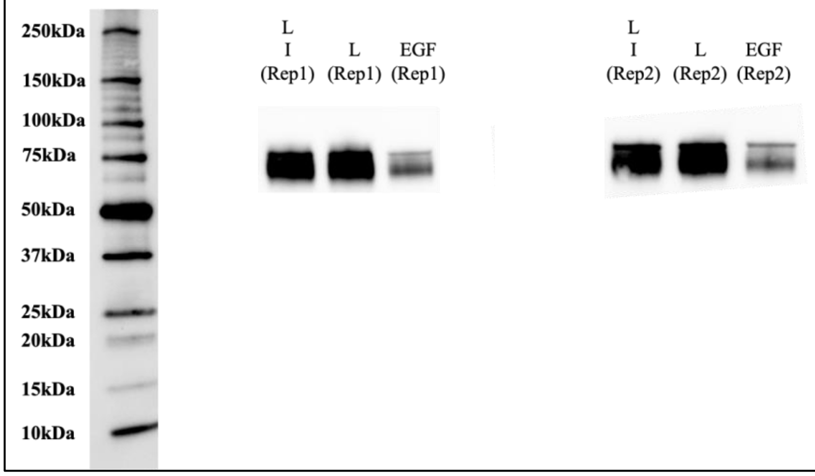


**Protein:** PDCD4/ Q53EL6, MW = 60 kDa, upregulated in L vs EGF and L/I vs EGF (cytoplasmic fraction)

**Conditions:**

- Primary Ab –PDCD4 (CST-9535) 1:1000
- Secondary Ab – Anti Rabbit HRP conjugate (CST- 7074S) 1:2000

*Note: L-lapatinib, I-ipatasertib*

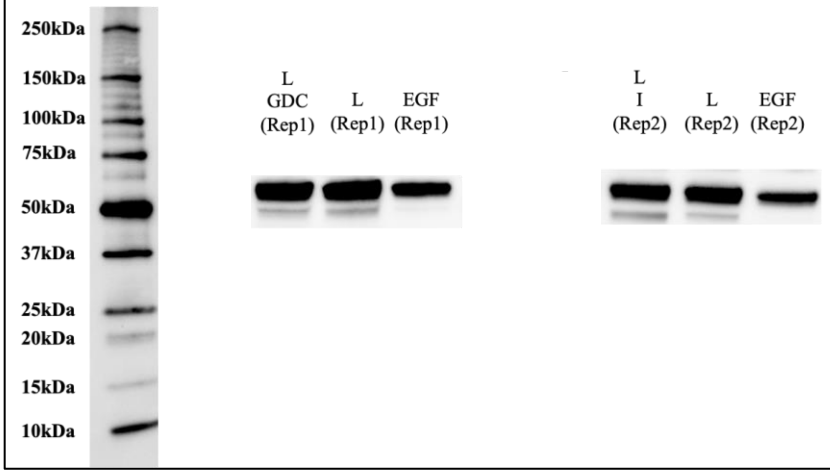

**Protein:** CD82/ P27701, MW = 30-90 kDa, upregulated in L vs EGF and L/I vs EGF (cytoplasmic fraction)
**Conditions:**

- Primary Ab –CD82 (CST-12439) 1:1000
- Secondary Ab – Anti Rabbit HRP conjugate (CST- 7074S) 1:2000

*Note: L-lapatinib, I-ipatasertib*

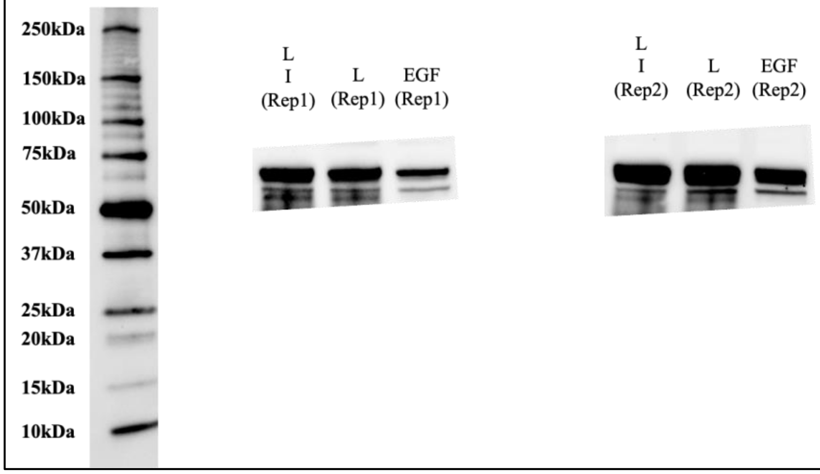


**Protein:** 14-3-3 sigma/ P31947, MW = 28 kDa, downregulated in L vs EGF and L/I vs EGF (cytoplasmic fraction)
**Conditions:**

- Primary Ab –14-3-3 sigma (RD-AF4424) 1:200
- Secondary Ab – Anti Goat HRP conjugate (RD- HAF017) 1:1000

*Note: L-lapatinib, I-ipatasertib*

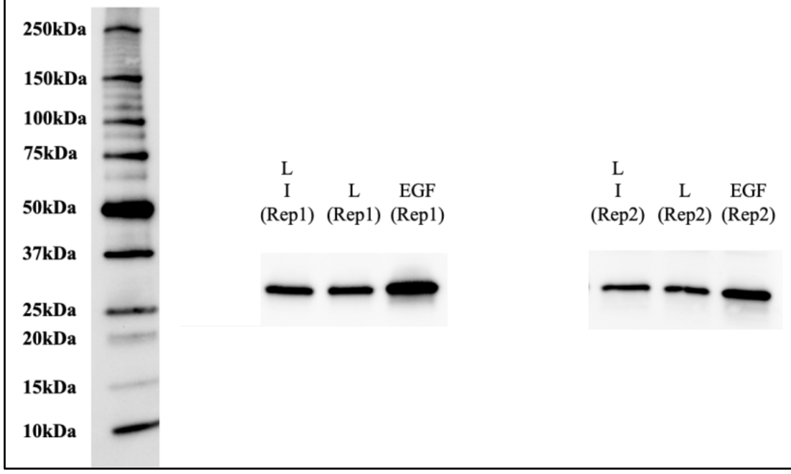

**Protein:** TOP2A/ P11388, MW = 190 kDa, downregulated in L vs EGF and L/I vs EGF (nuclear fraction)
**Conditions:**

- Primary Ab –TOP2A (CST-12286) 1:1000
- Secondary Ab – Anti Rabbit HRP conjugate (CST- 7074S) 1:2000

*Note: L-lapatinib, I-ipatasertib*

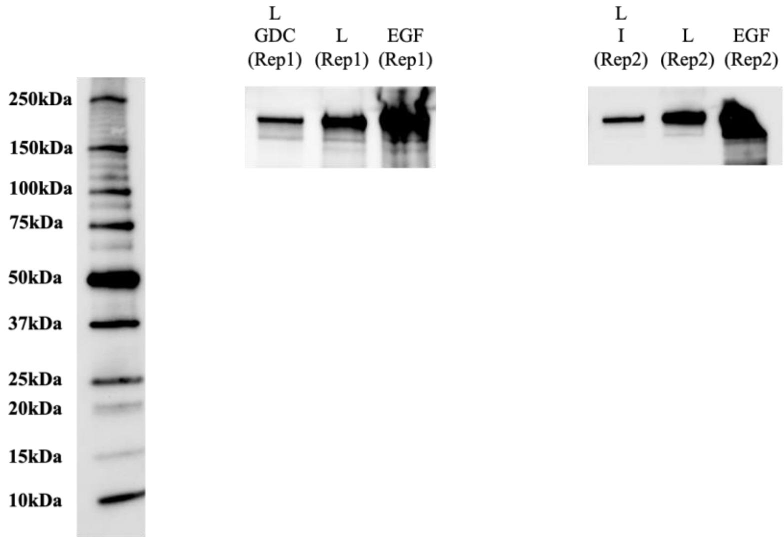
